# Amyotrophic Lateral Sclerosis-associated 3′ UTR enhancer embedded within *CAV1* risk gene

**DOI:** 10.1101/2025.09.09.675029

**Authors:** Laura J. Harrison, Tobias Moll, Johnathan Cooper-Knock, Daniel A. Bose

**Affiliations:** School of Biosciences, University of Sheffield, United Kingdom; Institute for Nucleic Acids, University of Sheffield, United Kingdom; Sheffield Institute for Translational Neuroscience, University of Sheffield, United Kingdom

**Keywords:** Enhancer, 3^′^ untranslated region (3^′^ UTR), intragenic enhancer, enhancer RNA (eRNA), amyotrophic lateral sclerosis (ALS), Caveolin-1 (CAV1), single nucleotide polymorphism (SNP)

## Abstract

Enhancer elements that reside within 3^*′*^ untranslated regions (UTRs) are an understudied phenomenon. Given the independent regulatory functions of enhancers and 3^*′*^ UTRs - enhancers governing pre-transcriptional control of gene expression and 3^*′*^ UTRs mediating post-transcriptional regulation of messenger RNA (mRNA) fate - 3^*′*^ UTR-associated enhancers may integrate these complementary layers to coordinate gene expression across multiple regulatory stages. Non-coding variation, impacting regulatory DNA, underpins the genetic architecture of disease. Indeed, the vast majority of single nucleotide polymorphisms (SNPs) associated with human complex diseases map to the non-coding genome, with causal variants particularly enriched within enhancers. Amyotrophic lateral sclerosis (ALS) is a complex neurodegenerative disorder associated with non-coding risk variants, many of which are increasingly linked to enhancer disruption. The *CAV1* gene, encoding the neuroprotective protein Caveolin-1, is a known ALS risk gene, yet the functional consequences of ALS-associated variation in its regulatory elements remain largely unexplored. Here, we combine genome-wide enhancer profiling with targeted experimental validation to define a previously uncharacterised ALS-associated enhancer embedded within the *CAV1* 3^*′*^ UTR, systematically assess its regulatory potential, and evaluate the impact of ALS-associated SNPs on enhancer function. We show that an individual ALS-associated SNP within this 3^*′*^ UTR-associated enhancer may disrupt function on multiple levels: at the DNA and chromatin level, by altering transcription factor binding with potential effects on recruitment of epigenetic co-regulators; and at the RNA level, by reshaping the structure and stability of a novel enhancer RNA transcribed from this locus. Collectively, our findings highlight this proximal *CAV1/CAV2* enhancer as a functionally important regulatory element embedded with the 3^*′*^ UTR of an ALS risk gene, illustrate how non-coding variants can impact multiple layers of gene regulation, and provide mechanistic insight into how intragenic enhancers contribute to ALS risk. More broadly, this work underscores the importance of 3^*′*^ UTR-associated enhancers as modulators of risk gene expression and underexplored contributors to human complex disease.

## Introduction

In contrast to single gene disorders, which are largely driven by protein-coding variants that often result in substantial phenotypic effects, complex diseases are caused primarily by non-coding variants which presumably affect gene regulation (Pickrell, 2014; Welter et al., 2014). In fact, the vast majority of single nucleotide polymorphisms (SNPs) associated with human complex diseases by genome wide association studies (GWAS) map to the non-coding genome, with causal disease variants enriched within *cis*-regulatory elements, particularly promoters and enhancers (Bal et al., 2022; ENCODE Project Consortium, 2012; Farh et al., 2015; Hindorff et al., 2009; van Arensbergen et al., 2019). Many complex trait alleles may therefore act by altering regulatory elements to influence gene regulation in a cell type-specific manner, specifically affecting cell types most relevant to disease phenotype (Farh et al., 2015; Finucane et al., 2015; Roadmap Epigenomics Consortium et al., 2015; Trynka et al., 2013). Yet, the mechanisms by which non-coding variants contribute to complex disease remain largely undefined.

Amyotrophic Lateral Sclerosis (ALS) is a late-onset progressive neurodegenerative disorder characterised by the degeneration of the upper and lower motor neurons, presenting with lower motor neuron-related muscle atrophy resulting from denervation, and upper neuron-related sclerosis of the corticospinal tract and its cortical origins (reviewed in Bäumer et al., 2014). The vast majority of ALS cases are considered to be sporadic, likely resulting from interactions between genetic and environmental factors in predisposed individuals, indicative of ALS as a complex disease (reviewed in Bäumer et al., 2014). Despite genetic factors being estimated to account for up to 52% of the variance in ALS development risk in a population-based parent-offspring heritability study, the majority of sporadic ALS cases have no associated genetic risk factor identified (Ryan et al., 2019). The bulk of ALS heritability is accounted for by SNPs with associations below genome-wide significance, potentially implying that many genetic risk variants of weak effect size are yet to be discovered and may account for the “missing heritability” associated with ALS as a complex disease (Manolio et al., 2009; Speed et al., 2012; van Rheenen et al., 2016). ALS-associated SNPs are increasingly linked to disruption in enhancer function (Cooper-Knock et al., 2021; Zhang et al., 2022; Yousefian-Jazi et al., 2020). Sequence polymorphisms within enhancer elements may contribute to ALS aetiology via alterations in transcription factor (TF) binding, chromatin architecture, or enhancer RNA (eRNA) structure and function (reviewed in Carullo and Day, 2019).

The *CAV1* gene, encoding the membrane lipid raft scaffold protein Caveolin-1 (CAV1), has been established as an ALS risk gene, with functional genetic variants enriched within the *CAV1* coding sequence in ALS patients (Cooper-Knock et al., 2021). CAV1 is a key protein component of caveolae, submicroscopic invaginations in the plasma membranes abundant in many mammalian cell types (reviewed in Parton and del Pozo, 2013). Converging evidence suggests a neuroprotective role for CAV1 in ALS (Cooper-Knock et al., 2021; Egawa et al., 2018; Head et al., 2011, 2008; Sawada et al., 2019; Takayasu et al., 2010). Indeed, *CAV1* (and *CAV2*) gene expression is consistently significantly higher in post-mortem brain tissues of ALS patients versus healthy control individuals (Adey et al., 2023). Beyond the functional link between *CAV1* coding variants and ALS, rare variant burden analysis has uncovered ALS-associated variants within enhancers linked with both the *CAV1* gene, and the paralogous Caveolin-2-encoding gene, *CAV2* (Cooper-Knock et al., 2021). Such pathogenic enhancer variants have been hypothesised to result in reduced transcription of their target genes, with the associated reduction in *CAV1* /*CAV2* expression proposed to induce neurotoxicity via the disruption of the heterooligomeric caveolin complex within membrane lipid rafts of motor neurons, resulting in impaired cell signalling (Cooper-Knock et al., 2021; Scherer et al., 1997; Head et al., 2008, 2010). However, the regulatory activity of candidate *CAV1/CAV2* enhancers containing ALS-associated variants has not been directly tested.

Here, we combine genome-wide enhancer profiling with targeted experimental validation of a previously uncharacterised proximal *CAV1/CAV2* enhancer within the *CAV1* 3^*′*^ UTR, systematically assessing its enhancer potential and examining how ALS-associated variants within this element may influence enhancer activity. This intragenic location represents a rare example of an ALS-associated enhancer embedded within the 3^*′*^ UTR of an established ALS risk gene, offering a unique opportunity to investigate how non-coding variants may modulate enhancer function while directly overlapping a disease-relevant host gene. Enhancer function is governed by a complex interplay of factors at multiple levels: at the chromatin level, through compartmentalisation (Dixon et al., 2012; Nora et al., 2012; Rao et al., 2014), histone modifications (Creyghton et al., 2010; Heintzman et al., 2007; Rada-Iglesias et al., 2011), and accessibility (Ernst et al., 2011); at the DNA level, through proximity to target promoters (Zuin et al., 2022) and the presence of TF binding motifs (reviewed in Spitz and Furlong, 2012); and at the transcriptional level, through the production and regulatory roles of eRNAs (De Santa et al., 2010; Hah et al., 2011; Kaikkonen et al., 2013; Kim et al., 2010; Li et al., 2013). This multi-dimensional complexity presents a major challenge in deciphering how enhancer variants contribute to complex disease. We hypothesise that ALS-associated SNPs within enhancer regions may disrupt one or multiple of these molecular features, thereby contributing to disease phenotype. Importantly, given the dual genic–regulatory context of 3^*′*^ UTR-associated enhancers, the effects of these variants may reflect disruption of enhancer activity, 3^*′*^ UTR function, or both. Understanding how such variants shape *CAV1* regulation therefore provides a broader framework for linking non-coding genetic risk to pathogenic mechanisms in ALS.

## Results

### Chromatin organisation and genomic location of ALS-associated candidate enhancers

Enhancers located in closer proximity to their target genes - be that in close linear genomic distance (Zuin et al., 2022) or through preferential interactions in 3D space (Dixon et al., 2012; Nora et al., 2012; Sexton et al., 2012; Shen et al., 2012) - are often associated with stronger regulatory activity. We examined whether such preferential enhancer-gene proximity is reflected in the chromatin organisation of the candidate *CAV1/CAV2* enhancer landscape in ALS patient-derived motor neurons.

In order to assess the 3D chromatin architecture of the candidate *CAV1/CAV2* enhancer landscape in the context of ALS, we utilised published Hi-C count matrices from ALS patient fibroblast-derived induced pluripotent stem cells (iPSCs) differentiated *in vitro* into motor neurons, downloaded from the ENCODE portal; accession numbers listed in Table 1 (ENCODE Project Consortium, 2012; Hitz et al., 2023; Luo et al., 2020). Topologically associating doamins (TADs) were assigned at a 100kb resolution, while sub-TAD structures were resolved at 10kb to capture finer-scale chromatin interactions (Figure 1a). While TAD organisation is largely invariant regardless of cell type (Dixon et al., 2012; Nora et al., 2012; Dixon et al., 2015), sub-TADs can reconfigure in a cell type-specific manner (Dowen et al., 2014; Ji et al., 2016; Phillips-Cremins et al., 2013; Rao et al., 2014). To assess fine-scale chromatin structure conservation, we identified whether CTCF in neuroblastoma model cell lines binds at regions identified as sub-TAD boundaries in *in vitro* differentiated motor neurons. Using published CTCF ChIP-seq data from SK-N-SH neuroblastoma cell lines, downloaded from the ENCODE portal, accession number listed in Table 1 (ENCODE Project Consortium, 2012; Hitz et al., 2023; Luo et al., 2020), we identified enrichment of CTCF ChIP-seq signal at sub-TAD boundaries (Figure 1a; Figure S1), consistent with stable boundary positioning between these two neurally-relevant cell types.

**Table 1.**
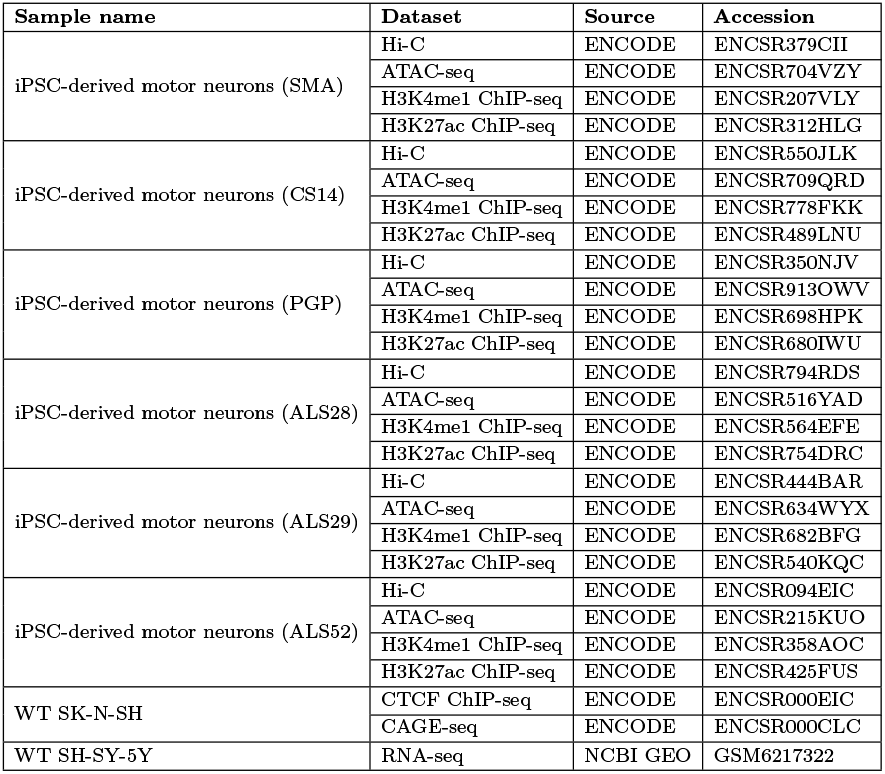
Sources of published datasets used in this study.

**Fig. 1:**
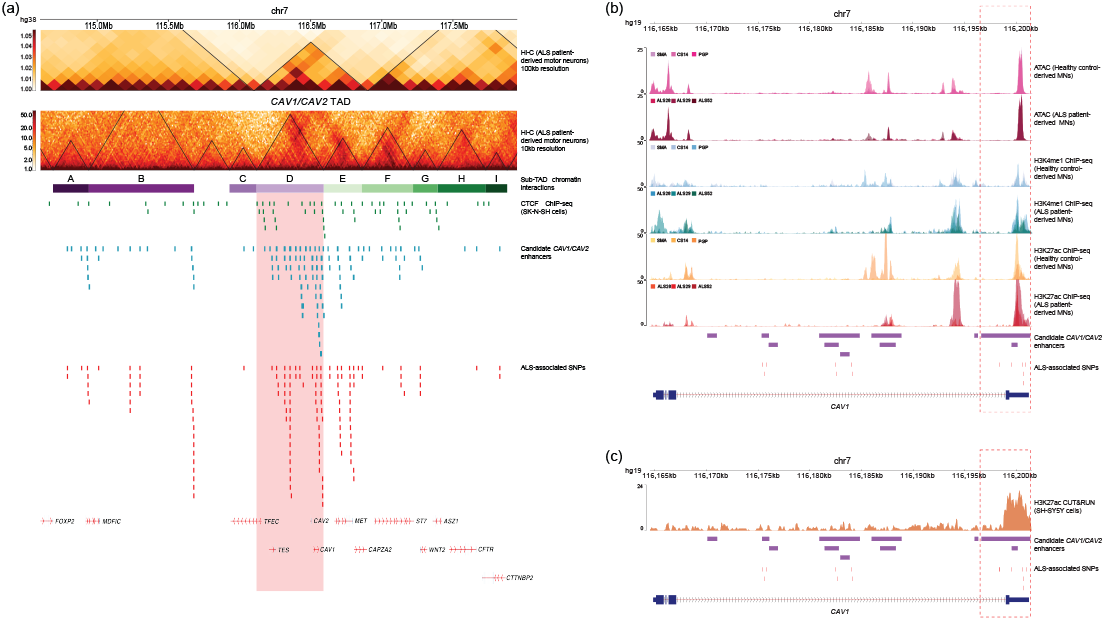
Chromatin organisation of ALS-associated *CAV1/CAV2* enhancers (a) Chromosome topology, as determined by Hi-C contact frequencies, at both 100kb resolution and 10kb resolution, at chr7:114,600,000-117,950,000 (hg38) in patient-derived iPSCs differentiated *in vitro* into motor neurons (Zhang et al., 2022); CTCF binding as determined by CTCF ChIP-seq in SK-N-SH neuroblastoma cells (ENCODE Project Consortium, 2012; Hitz et al., 2023; Luo et al., 2020); candidate *CAV1/CAV2* enhancers identified by the ABC model (Cooper-Knock et al., 2021; Fulco et al., 2019); ALS-associated SNPs within candidate *CAV1/CAV2* enhancers identified by GWAS (Cooper-Knock et al., 2021). (b) Trackline at chr7:116,164,500-116,201,500 (hg19) demonstrating ATAC-seq in patient-derived motor neurons; H3K4me1 ChIP-seq in patient-derived motor neurons; H3K27ac ChIP-seq in patient-derived motor neurons (Zhang et al., 2022); candidate *CAV1/CAV2* enhancers identified by the ABC model (Cooper-Knock et al., 2021; Fulco et al., 2019) and ALS-associated SNPs (Cooper-Knock et al., 2021). Position of the *CAV1/CAV2* proximal enhancer (chr7:116,196,622-116,201,351) is highlighted by red dashed box. (c) Trackline at chr7:116,164,500-116,201,500 (hg19) demonstrating H3K27ac CUT&RUN in SH-SY5Y cells; candidate *CAV1/CAV2* enhancers identified by the ABC model (Cooper-Knock et al., 2021; Fulco et al., 2019) and ALS-associated SNPs (Cooper-Knock et al., 2021). Position of the *CAV1/CAV2* proximal enhancer (chr7:116,196,622-116,201,351) is highlighted by red dashed box.

Both *CAV1* and *CAV2* were located within the same TAD (Figure 1a), which, despite its relatively small sub-megabase size compared to adjacent TADs, contained 64% of all candidate enhancers in our dataset, predicted to be associated with *CAV1/CAV2* by the Activity-By-Contact (ABC) model (Fulco et al., 2019; Cooper-Knock et al., 2021). Across the candidate enhancer landscape, nine sub-TADs (A-I) were identified (Figure 1a), with *CAV1* and *CAV2* residing in sub-TAD D. Notably, 48% of enhancers were located within sub-TAD D, supporting a preferential spatial association between *CAV1* and its putative enhancers in ALS patient-derived motor neurons.

To explore whether candidate enhancers in close spatial proximity to *CAV1/CAV2* harbour ALS-associated SNPs, we quantified ALS-associated SNPs – identified by GWAS as part of Project MiNE (Cooper-Knock et al., 2021) – within candidate *CAV1/CAV2* enhancers located in the same TAD as *CAV1* and *CAV2*. Consistent with the enrichment of candidate enhancers within the same TAD as *CAV1* and *CAV2*, 70% of the ALS-associated SNPs in our dataset were within enhancers located with the *CAV1/CAV2* TAD. Similarly, 48% of the ALS-associated SNPs resided within enhancers in sub-TAD D, mirroring the enrichment of candidate enhancers in this domain.

Taken together, we have demonstrated that both candidate *CAV1/CAV2* enhancers - identified via the ABC model (Fulco et al., 2019) - and ALS-associated SNPs within these enhancers - identified via GWAS and burden testing (Cooper-Knock et al., 2021) - preferentially reside within the same chromatin domain as the *CAV1* and *CAV2* genes. This pattern persists both at the level of higher-order chromatin organisation (TADs) and finer-scale chromatin structures (sub-TADs).

### Biochemical annotation of ALS-associated candidate *CAV1/CAV2* enhancers

Next, we characterised the chromatin signatures of candidate *CAV1/CAV2* enhancers to identify disease-relevant, putatively active enhancers. Specifically, we aimed to identify enhancers of interest that were in accessible chromatin and co-enriched for characteristic enhancer-associated histone modifications across ALS patient-derived motor neuron samples, enabling the selection of enhancers for subsequent functional annotation in SH-SY5Y cells. To explore chromatin accessibility at candidate *CAV1/CAV2* enhancers, we have utilised published ATAC-seq datasets from iPSC-derived motor neurons from healthy controls and ALS patients (Zhang et al., 2022), downloaded from the ENCODE portal; accession numbers listed in Table 1 (ENCODE Project Consortium, 2012; Hitz et al., 2023; Luo et al., 2020). To explore the distribution of enhancer-associated histone modifications, H3K4me1 and H3K27ac, across the candidate *CAV1/CAV2* enhancer landscape, we have utilised published ChIP-seq datasets from the same iPSC-derived motor neuron lines (Zhang et al., 2022), downloaded from the ENCODE portal; accession numbers listed in Table 1 (ENCODE Project Consortium, 2012; Hitz et al., 2023; Luo et al., 2020).

We identified a proximal enhancer region of interest, spanning chr7:116,196,622-116,201,351, that is located across the 3^*′*^ UTR of the *CAV1* gene, henceforth referred to as the proximal *CAV1/CAV2* enhancer. This enhancer was in accessible chromatin in all healthy control-derived and patient-derived *in vitro* differentiated motor neuron samples, as indicated by ATAC-seq signal enrichment (Figure 1b). Furthermore, this enhancer region was co-enriched for H3K4me1 and H3K27ac ChIP-seq signal in all ALS patient-derived *in vitro* differentiated motor neuron lines, and the eldest healthy control-derived sample (Figure 1b).

With the candidate proximal *CAV1/CAV2* enhancer displaying chromatin features characteristic of an active enhancer in ALS patient-derived motor neurons, we next aimed to functionally characterise this enhancer and assess the impact of ALS-associated SNPs in the SH-SY5Y neuroblastoma model cell line. Firstly, to confirm that this enhancer remains putatively active in SH-SY5Y cells, we utilised CUT&RUN for H3K27ac in SH-SY5Y cells to identify transcriptionally active enhancers in this model cell line. We observed a significant enrichment for H3K27ac CUT&RUN signal across the proximal enhancer region (Figure 1c). This enhancer region demonstrated the highest H3K27ac CUT&RUN signal enrichment of all of the candidate *CAV1/CAV2* enhancers in our dataset, with a 13.4-fold enrichment in H3K27ac signal over histone H3 signal. This suggests that, akin to in ALS patient-derived motor neurons, this candidate enhancer region is predicted to be active in SH-SY5Y cells.

### How does repression of ALS-associated enhancers impact *CAV1* expression?

Next, we aimed to functionally characterise ALS-associated *CAV1* /*CAV2* enhancer regions to validate enhancer-gene relationships between putative enhancer regions of interest and *CAV1*. Despite biochemical annotation data supporting enhancer identity for the proximal *CAV1/CAV2* enhancer region, this alone does not confirm a direct regulatory relationship with *CAV1*. To validate that the proximal *CAV1/CAV2* enhancer does indeed regulate *CAV1* expression, we targeted the CRISPR interference (CRISPRi) effector, dCas9-Zim3, to the proximal enhancer, and for comparison, to the candidate *CAV1* promoter and to a recently characterised distal *CAV1/CAV2* enhancer located *∼*20kb downstream (chr7:116,220,523-116,225,513), hereafter referred to as the distal *CAV1/CAV2* enhancer (Cooper-Knock et al., 2021), and observed the impact on *CAV1* expression (Figure 2a,b).

**Fig. 2:**
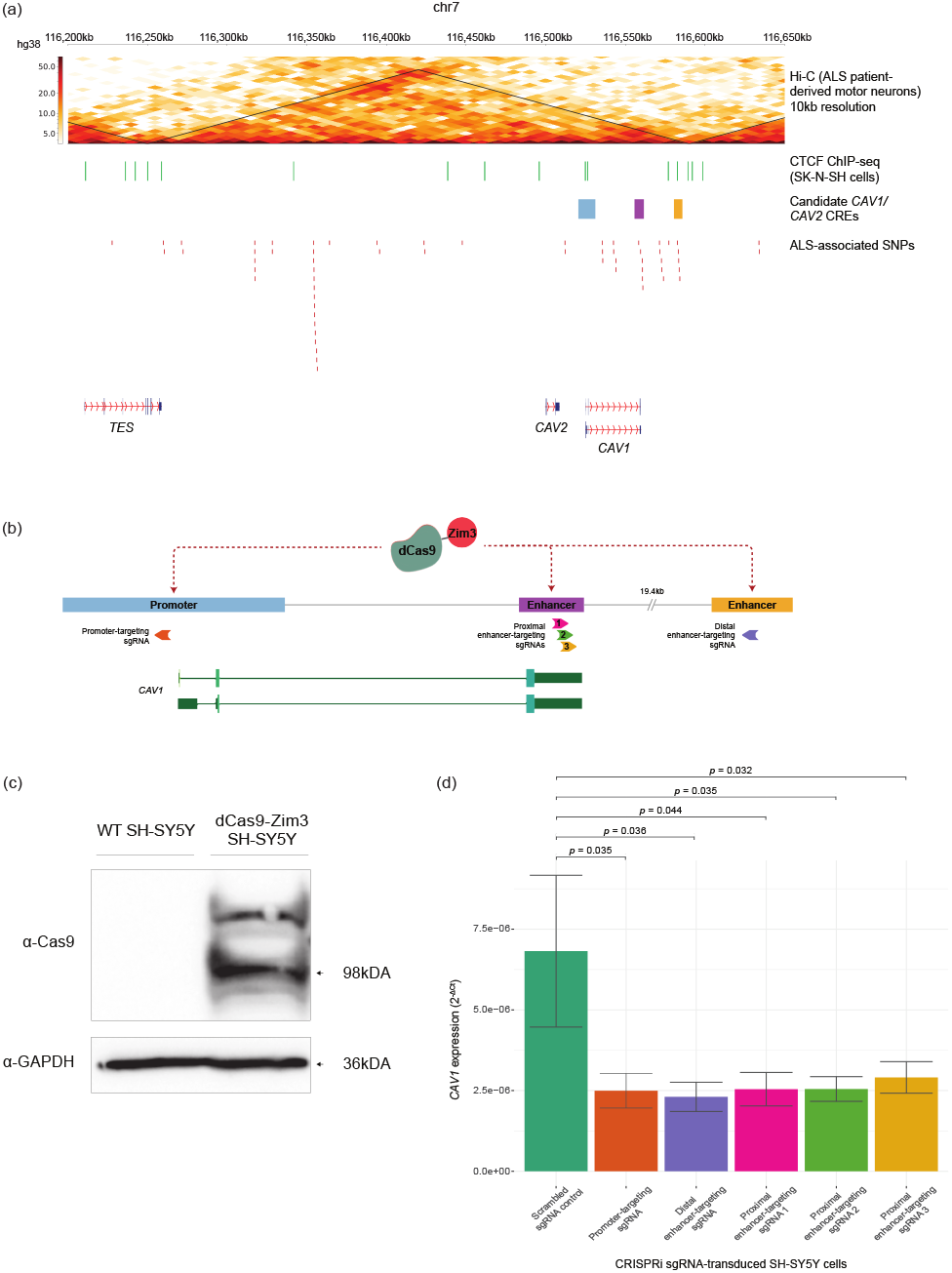
CRISPRi confirms identity of *CAV1* enhancers (a) Hi-C contact frequencies at chr7:116,200,000-116,650,000 (hg38) in patient fibroblast-derived iPSCs differentiated in vitro into motor neurons (Zhang et al., 2022), CTCF binding as determined by CTCF ChIP-seq in SK-N-SH neuroblastoma cells (ENCODE Project Consortium, 2012, Hitz et al., 2023, Luo et al., 2020), candidate *CAV1 cis*-regulatory elements identified by the ABC model (Cooper-Knock et al., 2021, Fulco et al., 2019) and ALS-associated SNP within candidate CAV1/CAV2 enhancers identified by GWAS (Cooper-Knock et al., 2021). Candidate *CAV1* promoter is shown in blue, proximal enhancer is shown in purple and distal enhancer is shown in yellow. (b) sgRNA target locations utilised for targeting of dCas9-Zim3 to candidate *CAV1* promoter and enhancers. (c) Western blot demonstrating stable expression of dCas9 in dCas9-Zim3 SH-SY5Y cells, confirmed by probing with *α*-Cas9 (Abcam, ab191468) and *α*-GAPDH (Proteintech, 60004-1-Ig). (d) Expression of *CAV1* mRNA in CRISPRi SH-SY5Y cell lines, as determined by RT-qPCR. Error bars determined by standard error, p-values determined by Student’s t-test. n=6

We generated a SH-SY5Y cell line stably expressing the dCas9-Zim3 CRISPRi effector (via transduction with pHR-UCOE-SFFV-Zim3-dCas9-P2A-Hygro), with Cas9 expression confirmed via western blot (Figure 2c). This cell line was further transduced with lentiGuide-Puro constructs containing sgRNAs targeting the regulatory regions of interest and *CAV1* pre-mRNA expression was subsequently measured via RT-qPCR. As expected of regulatory elements implicated in *CAV1* expression, targeting the CRISPRi effector to both the proximal and distal enhancers, in addition to the promoter, significantly reduced *CAV1* pre-mRNA expression compared to a scrambled control (Figure 2d). Notably, CRISPRi targeting of the distal enhancer resulted in a 66% reduction in *CAV1* expression relative to scrambled control, while targeting the proximal enhancer yielded a 57–63% reduction, depending on the sgRNA used. These reductions in expression are of a comparable magnitude to that observed when targeting the candidate promoter, which induced a 63% relative reduction in *CAV1* expression, supporting that both the proximal and distal enhancers act as bona fide *CAV1* enhancers.

### ALS-associated SNPs alter the regulatory function of the proximal *CAV1/CAV2* enhancer

To quantitatively evaluate the regulatory potential of the proximal *CAV1/CAV2* enhancer, we performed a luciferase reporter assay in SH-SY5Y cells, with luminescence produced by the reporter gene directly proportional to the capacity of the enhancer to drive mammalian gene expression *in vitro*. Beyond testing the regulatory function of the WT proximal *CAV1/CAV2* enhancer region, we also employed the luciferase reporter system to test the basal regulatory impact of ALS-associated SNPs on enhancer function. We introduced 5 ALS-associated SNPs into the enhancer sequence of the reporter construct: chr7:116,199,522T*>*C, chr7:116,200,589T*>*A, chr7:116,200,705C*>*T, chr7:116,200,719C*>*A and chr7:116,200,953A*>*T (Figure 3a). As controls, we included a negative control reporter construct lacking both the SV40 promoter and enhancer (Figure 3b) and a positive control construct containing both the SV40 promoter and enhancer (Figure 3c).

**Fig. 3:**
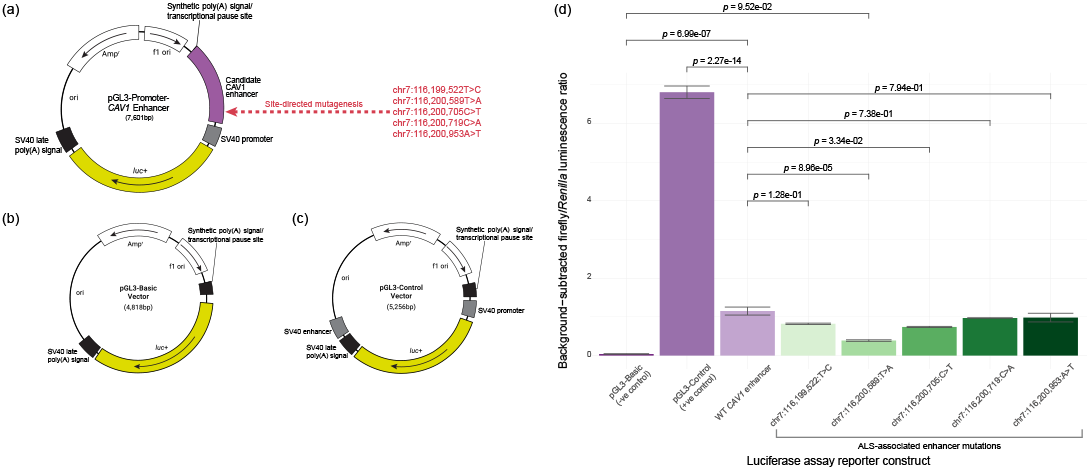
ALS-associated SNPs alter the regulatory function of the proximal *CAV1/CAV2* enhancer (a) Plasmid map of the luciferase reporter vector, modified from pGL3-Promoter, containing the proximal *CAV1/CAV2* enhancer with either the wild-type sequence or with ALS-associated SNPs introduced by site-directed mutagenesis. (b) Plasmid map of the pGL3-Basic negative control reporter vector. (c) Plasmid map of the pGL3-Control positive control reporter vector. (d) Background-subtracted firefly/*Renilla* luminescence ratio for the firefly reporter construct containing WT and mutated enhancer sequences in SH-SY5Y cells. Error bars determined by standard error, *p*-values determined by one-way ANOVA by Tukey’s Honestly Significant Difference (HSD) test. n = 3

While the proximal *CAV1/CAV2* enhancer appeared to be a comparatively weak enhancer, relative to the strong SV40 enhancer, it drove significantly more *Renilla* luciferase-normalised firefly luciferase expression than the negative control (Figure 3d). Furthermore, two ALS-associated mutations, chr7:116,200,589T*>*A and chr7:116,200,705C*>*T, significantly impaired enhancer-driven luciferase expression *in vitro*, compared to the WT enhancer sequence. Notably, the enhancer sequence containing the chr7:116,200,589T*>*A SNP failed to drive luciferase expression beyond the levels observed in the negative control, which lacked an enhancer altogether (Figure 3d).

### Transcription at candidate *CAV1/CAV2* enhancers

#### Transient transcriptome sequencing is capable of capturing transient non-coding transcripts

To detect eRNA transcription in SH-SY5Y cells we utilised transient transcriptome sequencing (TT-seq). TT-seq couples metabolic labelling with 4-thiouridine (4sU) with RNA fragmentation for the high-throughput capture of transient non-coding RNAs (ncRNAs) prior to degradation by the RNA exosome (Schwalb et al., 2016), and thus has great potential to capture eRNAs, despite their transience (Figure 4a). For normalisation purposes, we have utilised a 4-thiouracil (4tU)-labelled *Saccharomyces cerevisiae* RNA spike-in. To demonstrate the efficacy of the spike-in for controlling variation in global gene expression counts between replicates, we compared the gene counts between unnormalised versus spike-in normalised TT-seq data (Figure 4b). We observed a very strong positive linear relationship between the gene expression counts of the two replicates of WT SH-SY5Y TT-seq (Figure 4c).

**Fig. 4:**
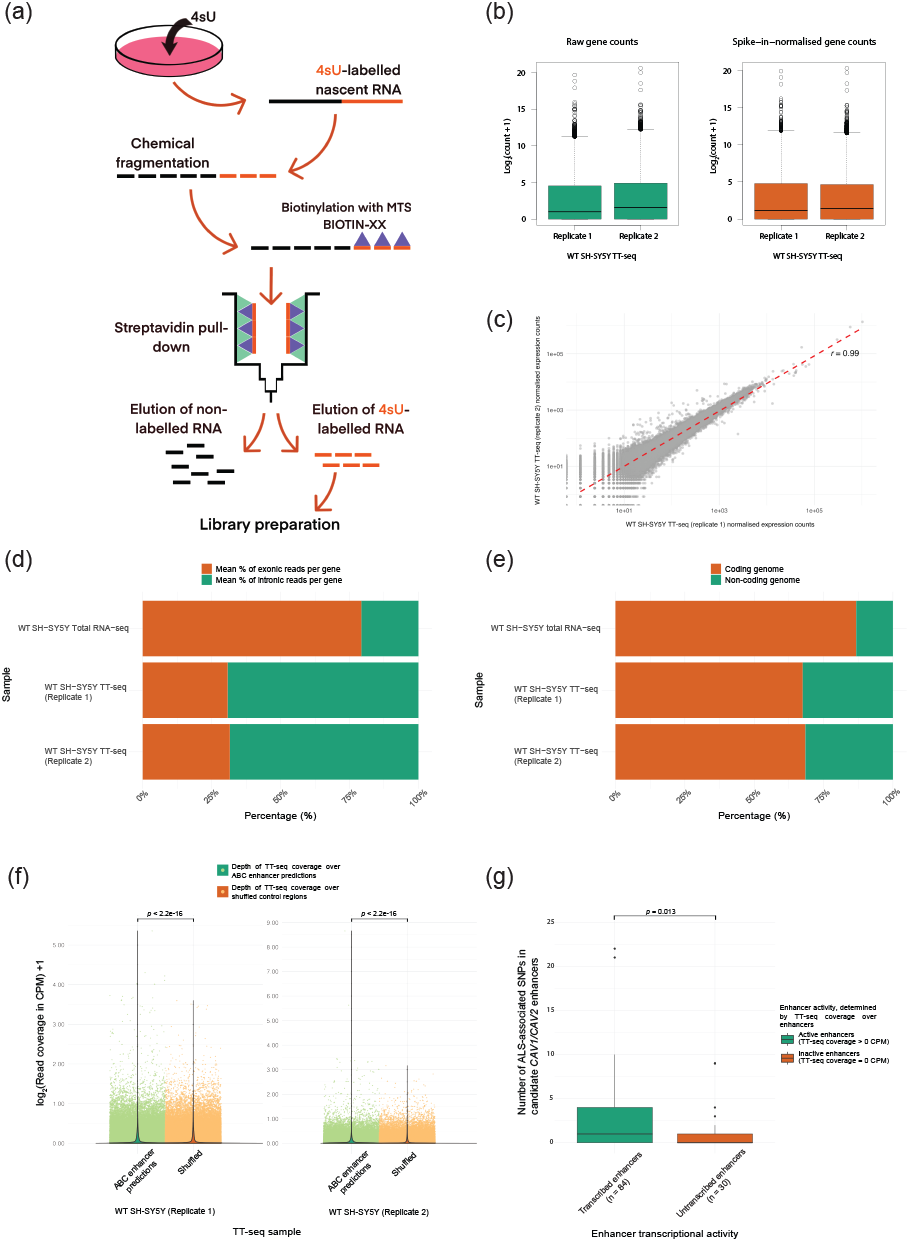
Transient transcriptome sequencing can capture eRNAs genome-wide (a) Schematic representing the workflow of TT-seq. Nascent RNA is metabolically labelled *in vivo* via nucleoside analogue, 4-thiouridine (4sU). Total RNA is extracted and fragmented. 4sU-labelled fragmented RNA is biotinylated and captured using steptavidin pull-down prior to library preparation and sequencing (Gregersen et al., 2020; Schwalb et al., 2016). (b) Log-transformed gene counts of WT SH-SY5Y TT-seq data from two replicates both prior and post spike-in normalisation. (c) Correlation of spike-in normalised gene counts between WT SH-SY5Y TT-seq replicates. Correlation strength quantified by the Pearson correlation coefficient. (d) Exonic to intronic per gene transcript ratios for TT-seq in SH-SY5Y cells, compared to total RNA-seq in SH-SY5Y cells (Liu et al., 2022), to recapitulate the expected values for TT-seq, with the coverage of intronic to exonic transcripts per gene estimated to be *∼*60% (Schwalb et al., 2016). (e) Proportion of total TT-seq reads in SH-SY5Y cells that map to non-coding regions of the genome, compared to the corresponding proportions observed in total RNA-seq data from the same cell type. (f) WT SH-SY5Y TT-seq coverage (*log*_2_ CPM+1) over ABC candidate enhancers (Fulco et al., 2019) compared to shuffled control regions. *p*-values determined by Wilcoxon Rank Sum. (g) Number of ALS-associated SNPs (Cooper-Knock et al., 2021) transcribed versus untranscribed candidate *CAV1/CAV2* enhancers (Cooper-Knock et al., 2021; Fulco et al., 2019), as determined by TT-seq in WT SH-SY5Y cells. *p*-value deteremined by Wilcoxon Rank Sum.

The coverage of transient intronic transcripts with respect to exonic transcripts is estimated to be *∼*60% for TT-seq, compared to *∼*8% for total RNA-seq (Schwalb et al., 2016). To validate the usage of TT-seq in SH-SY5Y cells for the capture of transient non-coding transcripts, we investigated whether we could recapitulate this ratio in our samples. For each given GENCODE gene annotation (Frankish et al., 2019), the ratio of intronic versus exonic transcripts was calculated, for both our TT-seq data and a published total RNA-seq dataset from WT SH-SY5Y cells (Liu et al., 2022), data accessible at NCBI GEO database (Edgar et al., 2002), accession number listed in Table 1. For TT-seq, we observed a mean percentage intron read count of *∼*69% and *∼*68% per gene, for replicate 1 and 2, respectively, compared to *∼*21% for total RNA-seq (Figure 4d). In order to identify if the capacity to capture unstable transcripts extended beyond introns, we investigated the TT-seq coverage over the non-coding genome as a whole. In comparison to total RNA-seq, where only 13.2% of the total reads mapped to the non-coding genome, 32.5% and 31.5% of the total reads mapped to non-coding genome for TT-seq replicate 1 and 2, respectively (Figure 4e).

#### Using nascent transcription data to identify active enhancers

Further, we explored the capacity for TT-seq to capture eRNAs in SH-SY5Y cells by investigating the TT-seq coverage over enhancer predictions from the ABC model (Fulco et al., 2019). As a control, we generated chromosome-matched, non-exonic shuffled regions by randomly permuting the genomic locations of the ABC enhancer predictions along the same chromosome, excluding coding exons. In a direct comparison of TT-seq read coverage between ABC enhancer regions and these non-exonic shuffled control regions, we observed significantly higher transcriptional activity at ABC enhancers (Figure 4f), consistent with TT-seq capturing eRNA transcription genome-wide.

To test whether disease-associated SNPs localise to active enhancers in physiologically relevant cell types, we investigated the number of ALS-associated SNPs in actively transcribed candidate *CAV1/CAV2* enhancers versus those absent of TT-seq read coverage. As expected, transcribed candidate *CAV1/CAV2* enhancers, as determined by TT-seq coverage, contain significantly more ALS-associated SNPs than untranscribed enhancers (Figure 4g). Transcribed enhancers contained on average 3 SNPs, whereas untranscribed enhancers contained on average 1 SNP, to the closest integer. Taken together, these results demonstrate the utility of TT-seq for capturing transient non-coding RNA species, including eRNAs transcribed from active enhancers, in SH-SY5Y cells. Furthermore, the enrichment of ALS-associated SNPs within transcribed enhancers, as detected by TT-seq highlights the biological relevance of this approach for linking enhancer activity to disease-associated genetic variation.

### ALS-associated SNPs are predicted to alter eRNA structure and stability

#### Identification of novel eRNAs in SH-SY5Y cells

While TT-seq effectively captures eRNA transcription, providing valuable insights into enhancer activity (Schwalb et al., 2016), it also detects nascent transcription of other RNA species, including pre-mRNAs and long non-coding RNAs (lncRNAs).

This broader transcriptional capture introduces potential biases when quantifying enhancer transcription, as signal attributed to eRNA production may be confounded by overlapping or nearby transcription from non-enhancer sources.

To mitigate this, we performed *de novo* transcript assembly from TT-seq data, comparing assembled transcripts to the reference transcriptome and excluding those that exactly matched or were fully contained within the reference annotation. We further filtered for transcripts both originating from and fully contained within candidate *CAV1/CAV2* enhancer loci, retaining only those detected in both SH-SY5Y TT-seq replicates. We identified a novel antisense transcript transcribed from the proximal *CAV1/CAV2* enhancer at chr7:116,196,623-116,201,351, henceforth referred to as *eCAV1_WT*, consistently assembled from both replicates of TT-seq (Figure 5a).

**Fig. 5:**
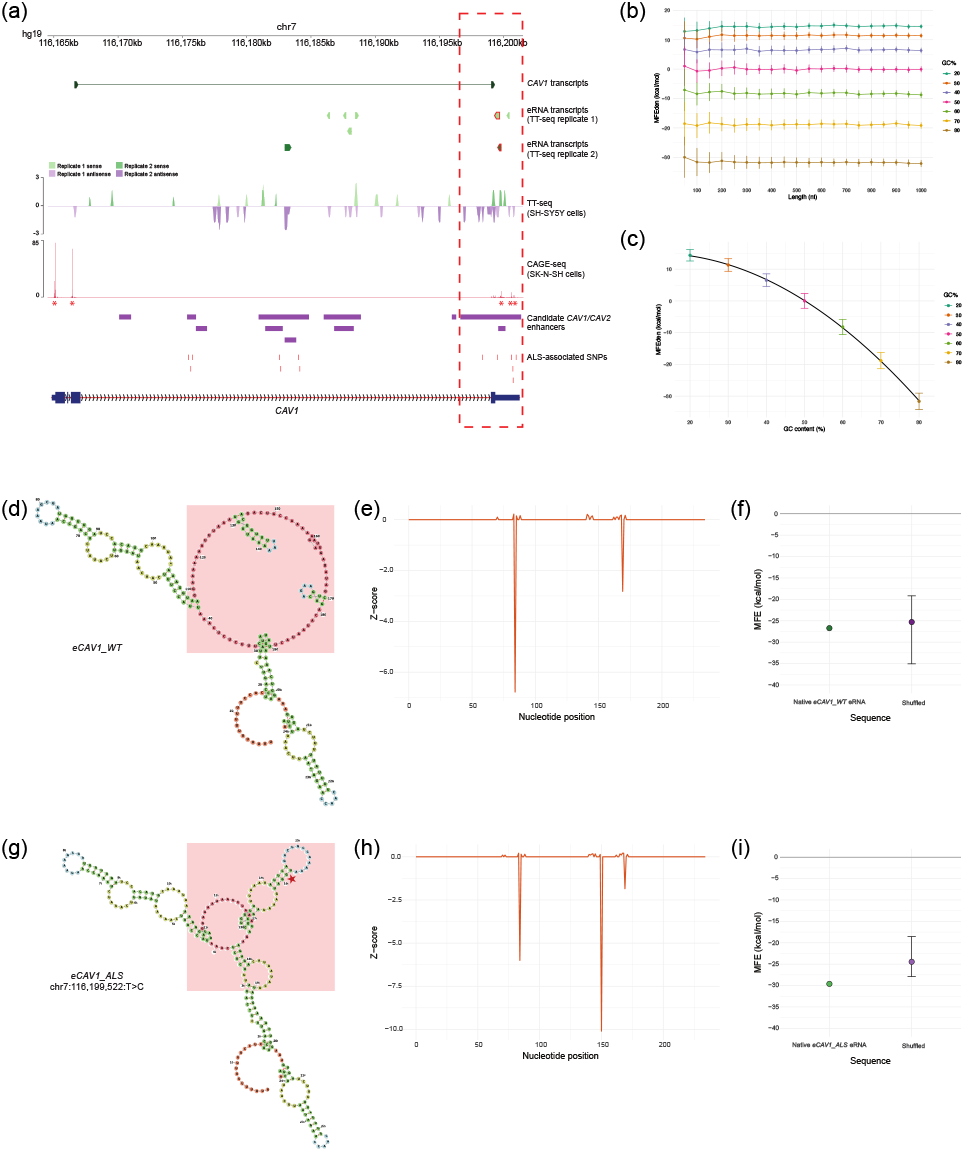
ALS-associated SNPs in the proximal *CAV1/CAV2* enhancer are predicted to alter eRNA structure and stability (a) Trackline at chr7:116,164,500-116,201,500 (hg19) demonstrating transcripts assembled from TT-seq data in WT SH-SY5Y cells, including the canonical *CAV1* mRNA transcript and novel *de novo* transcripts of enhancer origin and the corresponding TT-seq coverage; CAGE-seq in SK-N-SH neuroblastoma cells (ENCODE Project Consortium, 2012; Hitz et al., 2023; Luo et al., 2020); the candidate enhancers identified by the ABC model (Cooper-Knock et al., 2021; Fulco et al., 2019) and ALS-associated SNPs (Cooper-Knock et al., 2021). *eCAV1_WT* transcripts are highlighted with a red border. Transcription start sites, as determined by CAGE-seq peaks, denoted by red asterisks. (b) Mean MFEden versus RNA sequence length for shuffled sequences 50-1000 nt in length (in 50nt intervals) with GC content of 20%, 30%, 40%, 50%, 60%, 70% and 80%. Each point corresponds to the mean MFEden of 100 shuffled sequences. (c) Mean MFEden for sequences 50 nt to 1000 nt versus GC content. Line is fitted using a quadratic linear regression model. (d) MFE structure prediction for the novel *CAV1* enhancer-associated antisense eRNA *eCAV1_WT* (chr7:116,199,437-116,199,680). (e) Thermodynamic z-score across the length of *eCAV1_WT*. (f) Native *eCAV1_WT* eRNA MFE compared to MFE values of shuffled RNA sequences (n=10). (g) MFE structure prediction for the *CAV1* enhancer-associated antisense eRNA upon computational induction of the chr7:116,199,522T*>*C ALS-associated SNP, *eCAV1_ALS*. Location of SNP is denoted by the red star. SNP-dependent structural changes are highlighted with red boxes. (h) Thermodynamic z-score across the length of *eCAV1_ALS*. (i) *eCAV1_ALS* eRNA MFE compared to MFE values of shuffled RNA sequences (n=10).

To corroborate eRNA production from this enhancer - which overlaps the *CAV1* 3^*′*^ UTR - and confirm the presence of transcription start sites (TSSs) independent of genic transcription, we analysed publicly available Cap Analysis of Gene Expression and Sequencing (CAGE-seq) data from WT SK-N-SH cells, downloaded from the ENCODE portal, accession number listed in Table 1. (ENCODE Project Consortium, 2012; Hitz et al., 2023; Luo et al., 2020). CAGE-seq specifically captures the 5^*′*^ *m*^7^*G*-cap of RNAs (Shiraki et al., 2003), making it a robust method for detecting capped RNAs such as eRNAs, which like mRNAs, are often 5^*′*^-capped (Kristjánsdóttir et al., 2020). We observed complete overlap between the *eCAV1_WT* transcript and a CAGE-seq peak within the proximal *CAV1/CAV2* enhancer, supporting enhancer-derived transcription.

#### eRNA structure and stability prediction

RNA folding into secondary structures can be conceptualised as the traversal of a complex free energy landscape, where different secondary structures correspond to local energy minima. The minimum free energy (MFE) structure represents the most thermodynamically stable conformation under specific temperature and ionic conditions (Zuker and Stiegler, 1981). Dynamic programming algorithms, such as RNAfold from the ViennaRNA package (Lorenz et al., 2011; Hofacker et al., 1994) and RNAstructure (Reuter and Mathews, 2010), predict RNA secondary structure by iteratively evaluating the thermodynamic costs of possible intra-sequence base-pairing interactions to identify the globally minimal free energy structure (Zuker and Stiegler, 1981; McCaskill, 1990). Functional RNAs, including mRNAs and microRNA (miRNA) precursors, exhibit more negative MFE values than expected based on their nucleotide composition alone (Bonnet et al., 2004; Seffens and Digby, 1999), establishing RNA MFE as an indicator of not only potential RNA stability, but also functionality.

To assess the predicted stability and potential functionality of *CAV1/CAV2* enhancer-associated eRNAs, we performed RNA secondary structure prediction using RNAfold (Lorenz et al., 2011). To normalise predicted MFE values, we calculated the MFE density (MFEden) index, successfully excluding the predominant influence of sequence length on MFE, allowing for evaluation of the contributions of nucleotide composition and order to RNA structural stability, while reducing length-related bias (Trotta, 2014). The MFE (*MFE*) of a given RNA sequence with a nucleotide length (*L*) is normalised by calculating the MFEden using the following formula (Trotta, 2014):

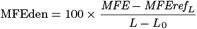

Where the *MFEref*_*L*_ is the expected MFE for an RNA sequence of *L* nucleotides with equimolar ratios of each nucleotide (A, C, G, and U) and *L*_0_ for MFEs computed by RNAfold is empirically equal to 8 nt (Trotta, 2014).

MFEden mitigates length dependence but remains sensitive to nucleotide composition and order, providing an estimate of how sequence features contribute to RNA structural stability independently of length (Trotta, 2014). Notably, pre-miRNAs typically exhibit lower MFEden values than expected based on nucleotide content alone (Trotta, 2014), highlighting the influence of nucleotide order on RNA folding in this functional RNA species. To separate the contributions of overall GC content from explicit nucleotide order to eRNA free energy values, we modelled expected MFEden as a function of GC content. We computed the MFEden of shuffled sequences ranging in length from 50 nt to 1000 nt (in 50 nt intervals) with GC contents of 20%, 40%, 50%, 60%, 70% and 80%, with 100 shuffled sequences generated for each iteration. Across lengths, GC content emerged as the dominant contributor to MFEden, remaining largely uniform for each given GC% (Figure 5b). We fit a second-degree polynomial regression to describe this non-linear relationship (Figure 5c), achieving an *R*^2^ of 0.979 on the training data and 0.978 on an independent test set. Residuals were centred near zero (*ē*_train_ = 1.90e-13; *ē*_test_ = 1.36e-01) with negligible correlation to GC content (*r*_train_ = -7.81e-13; *r*_test_ = -2.11e-03), indicating minimal bias (Figure S2).

In addition to MFEden, we employed the thermodynamic z-score to statistically evaluate the contribution of nucleotide order to eRNA stability (Andrews et al., 2018, 2022). Functional ncRNAs typically have significantly lower MFE values than randomised sequences with identical nucleotide compositions, reflecting ordered nucleotide arrangements forming stable local secondary structures (Babak et al., 2007; Clote et al., 2005).

The thermodynamic z-score compares the MFE of the native sequence to the mean MFE of multiple shuffled sequences, normalised by the standard deviation of all MFE values (Andrews et al., 2018, 2022; Clote et al., 2005):

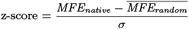

Where the *MFE*_*native*_ is the MFE of the native RNA sequence, *MFE*_*random*_ is the mean MFE of the shuffled RNA sequence and *σ* is the standard deviation of all MFE values. Negative z-scores indicate that the native sequence is more stable than expected by chance, isolating the effect of nucleotide order.

To further assess the contribution of nucleotide order to predicted RNA stability, we also compared the MFE of native eRNAs to that of 10 shuffled sequences of identical nucleotide composition and length, providing a benchmark to distinguish stability due to specific structural motifs from that driven by general nucleotide composition. This approach assumes that explicitly stable RNA motifs contribute more substantially to overall stability than base composition alone.

Together, these metrics provide complementary perspectives on RNA structural stability. Consistent results across them strengthen confidence in predicted eRNA stability, while discrepancies may reveal biases in structural determinants.

#### eRNA structure and stability prediction to model ALS-associated SNPs

To investigate whether ALS-associated SNPs possess the capacity to alter eRNA structure and stability, we focussed on the *eCAV1_WT* transcript, which overlaps the ALS-associated SNP chr7:116,199,522T*>*C within the proximal *CAV1/CAV2* enhancer. We computationally introduced this SNP into the *eCAV1_WT* sequence, generating a variant hereafter referred to as *eCAV1_ALS*. By comparing the predicted secondary structure of *eCAV1_ALS* to that of the native WT sequence, we assessed the potential structural and stability changes resulting from this single disease-associated nucleotide substitution.

We predicted the secondary RNA structure of *eCAV1_WT* (Figure 5d) using RNAfold, yielding an MFE of −26.73 kcal*/*mol and an observed MFEden (*MFEden*_*obs*_) of 12.69kcal/mol. Employing our polynomial prediction model to generate the expected MFEden (*MFEden*_*exp*_) for a sequence with the same GC content (28.28%) yielded the relatively more negative MFEden value of 12.06kcal/mol, suggesting that *eCAV1_WT* may be somewhat less stable than expected from nucleotide composition alone. The thermodynamic z-score across the eRNA averaged −0.032 with a minimum z-score of −6.78 (Figure 5e). Comparison of the MFE of *eCAV1_WT* to that of 10 shuffled sequences of identical nucleotide composition and length resulted in very similar MFE values (Figure 5e), indicating a limited contribution of specific local sequence motifs to overall RNA stability.

To model the impact of the chr7:116,199,522T*>*C SNP, we predicted the secondary RNA structure of *eCAV1_ALS* (Figure 5g), yielding a relatively more negative MFE of −29.61 kcal*/*mol and a slightly lower *MFEden*_*obs*_ of 11.47kcal/mol. Introduction of the SNP marginally increased the GC-content of the eRNA (28.69%), with the polynomial prediction model generating a *MFEden*_*exp*_ of 11.92kcal/mol. In contrast to *eCAV1_WT*, the *MFEden*_*obs*_ for *eCAV1_ALS* was slightly more negative than expected, suggesting that nucleotide order may enhance stability beyond the effect of GC content alone. In addition to the changes in the MFE, we visually observed alterations in the secondary RNA structure upon induction of the ALS-associated SNP. The large multiloop present in *eCAV1_WT* - with two small hairpin loops and a helix - was replaced in the *eCAV1_ALS* variant by a smaller multiloop containing two helices and an interior loop, in addition to a larger hairpin loop branching off the multiloop. In concordance with the reduction in MFE upon induction of the mutation, the mean thermodynamic z-score decreased to −0.066, with a minimum z-score of −10.1 (Figure 5h), indicating increased structural stability associated with the SNP. Of note, the minimum z-score in the WT eRNA sequence, and hence the most stable structure within the eRNA, corresponded to the stem loop 84 nt into the RNA sequence, whereas in the ALS variant it was attributed to a larger stem loop absent in the WT eRNA structure. Comparing the MFE of *eCAV1_ALS* to shuffled sequences revealed that the ALS SNP-containing eRNA had a consistently more negative MFE than random sequences (Figure 5i), indicating that, when compared to the WT sequence, introduction of the ALS-associated SNP enhanced structural stability.

### The predicted impact of ALS-associated SNPs on transcription factor binding and downstream coactivator function

#### The impact of CBP/p300 inhibition on *CAV1* mRNA and eRNA transcription

As previously established, the proximal *CAV1/CAV2* enhancer exhibits strong enrichment of H3K27ac CUT&RUN signal (Figure 1c), overlapping sites of *de novo* eRNA transcription (Figure 5a). CREB-binding protein (CBP)/p300 are key histone acetyltransferases responsible for depositing acetyl groups on histones flanking enhancers (Hilton et al., 2015; Narita et al., 2021; Miao et al., 2022). To determine whether CBP/p300 catalytic activity contributes to the regulation of *CAV1* expression and enhancer activity, we treated SH-SY5Y cells with A-485, a potent CBP/p300 catalytic inhibitor (Lasko et al., 2017), and quantified *CAV1* pre-mRNA and bidirectional eRNA expression at the proximal *CAV1/CAV2* enhancer by RT-qPCR relative to untreated cells (Figure 6a). A-485 treatment resulted in a reduction in overall *CAV1* pre-mRNA expression, relative to untreated SH-SY5Y cells. Indicative of CBP/p300-dependent activity at the proximal *CAV1/CAV2* enhancer, we also detected a significant decrease in both sense and antisense eRNA transcription upon A-485 treatment, compared to untreated SH-SY5Y cells (Figure 6b).

**Fig. 6:**
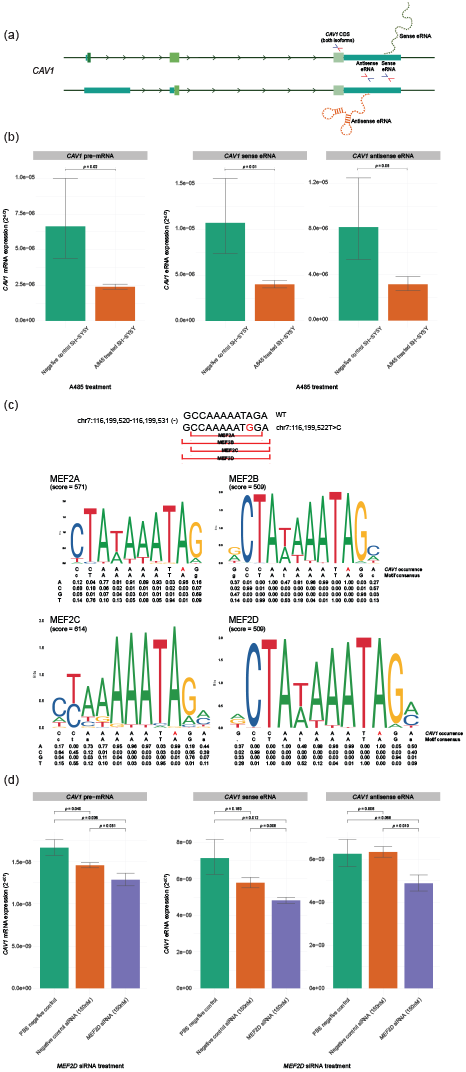
ALS-associated SNPs are predicted to impact transcription factor and cofactor binding at the proximal *CAV1/CAV2* enhancer (a) Schematic representation of the primers used for RT-qPCR. Sense and antisense eRNA primers designed based on assembled TT-seq transcripts. (b) RT-qPCR for overall *CAV1* pre-mRNA, sense and antisense eRNA expression in SH-SH5Y cells treated with A-485 compared to untreated SH-SY5Y cells. Error bars determined by standard error, *p*-values determined by unpaired t-test. n=3. (c) MEF2 binding site motifs and position weight matrices from the JASPAR database (Rauluseviciute et al., 2024) at chr7:116,199,520-116,199,531 (-), comparing the *CAV1* sequence occurrence and the motif consensus, with sites of the chr7:116,199,522T*>*C shown in red. (d) RT-qPCR for overall *CAV1* pre-mRNA, sense and antisense eRNA expression in SH-SH5Y cells treated with PBS (negative control), negative control siRNA and with *Mef2d* siRNA. Error bars determined by standard error, *p*-values determined by unpaired t-test. n=3.

#### ALS-associated SNPs are predicted to alter protein-binding motifs in *CAV1/CAV2* enhancers

Utilising the JASPAR database for TF binding profiles (Rauluseviciute et al., 2024), we investigated whether ALS-associated SNPs within the proximal *CAV1/CAV2* enhancer disrupt putative TF binding sites. After filtering for statistical significance (*p* = 10^−4^), we identified motifs for MEF2A, MEF2B, MEF2C and MEF2D that overlap with the site of the ALS-associated SNP chr7:116,199,522T*>*C. The MEF2 family was of particular interest, as MEF2 TFs are implicated in neuronal survival (Gong et al., 2003; Mao et al., 1999; Yang et al., 2009) and are key regulators of activity-dependent synapse development (Flavell et al., 2006; Shalizi et al., 2006). Homo- and heterodimers of MEF2 proteins bind directly to DNA regions possessing the MEF2 response element consensus sequence YTA(A/T)4TAR (Gossett et al., 1989), and nearly exact matches to this sequence were present on the reverse strand at chr7:116,199,520-116,199,531, placing the SNP directly within the binding motif (Figure 6c).

Position weight matrix (PWM) analysis revealed that at chr7:116,199,522, MEF2B and MEF2D have maximal binding preference for A, with a frequency of 1.00, suggesting that the chr7:116,199,522T*>*C SNP could strongly disrupt binding. MEF2A and MEF2C also exhibited strong preference for A at the same position, with frequencies of 0.96 and 0.99, respectively. With the T*>*C substitution resulting in a base change to a G on the reverse strand, binding potential for MEF2A and MEF2C may be retained, though diminished, with frequencies of 0.03 and 0.01 for G, respectively.

To test the functional impact of impaired MEF2 binding at the proximal *CAV1/CAV2* enhancer, we performed siRNA-mediated knockdown of *Mef2d* in SH-SY5Y cells and quantified *CAV1* mRNA and eRNA expression by RT-qPCR (Figure 6d). While we observed no significant difference in *CAV1* pre-mRNA expression between *Mef2d* siRNA-treated cells and control siRNA-treated cells, we observed a significant reduction in expression of both the sense and antisense *CAV1* eRNAs in *Mef2d* siRNA-treated cells, in comparison to control siRNA-treated cells.

Taken together, these results suggest that CBP/p300 and MEF2D activity is required for robust transcriptional activity at the proximal *CAV1/CAV2* enhancer, with the catalytic activity of CBP/p300 additionally required to support *CAV1* mRNA transcription.

## Discussion

We confirm the identity of an ALS-associated enhancer located within the 3^*′*^ UTR of the ALS risk gene *CAV1*. Prioritising enhancers for functional assessment based on their proximity to the *CAV1* promoter - under the assumption that closer enhancers may exert stronger regulatory effects (Zuin et al., 2022) - led to the identification of this 3^*′*^ UTR-associated enhancer. Within the *CAV1/CAV2* regulatory network, ALS-associated enhancer variants converge on a spatially clustered enhancer landscape, with both candidate CAV1/CAV2 enhancers and their embedded disease-associated SNPs preferentially residing within the same TAD and sub-TAD as *CAV1/CAV2* in ALS patient-derived motor neurons (Figure 1a). To our knowledge, this represents one of the first examples of a disease-associated enhancer located within the 3^*′*^ UTR of its own risk gene. This finding highlights a rare instance of an intragenic, disease-associated enhancer within an established ALS risk gene.

3^*′*^ UTR-associated enhancers remain an underexplored regulatory phenomenon, largely due to the challenges of distinguishing eRNA transcription from readthrough of the host gene into its 3^*′*^ UTR. For this reason, most enhancer discovery efforts have focused on intergenic and intronic regions, where the signatures of enhancer activity are less confounded by overlapping gene transcription. This challenge is exacerbated by the increasing recognition of transcription at the 3^*′*^ termini of protein-coding genes, which gives rise to terminus-associated non-coding RNAs (TANRs) with regulatory functions (Ni et al., 2020). At the *CAV1* 3^*′*^ UTR, biochemical profiling of enhancer-associated histone modifications indicated strong enhancer activity (Figure 1b, c), which we validated *in vitro* by CRISPRi and luciferase reporter assays (Figure 2, Figure 3). Profiling nascent transcription genome-wide using TT-seq in SH-SY5Y cells further revealed nascent transcription across the *CAV1* 3^*′*^ UTR (Figure 5a). To determine whether this observed nascent transcription represented eRNA rather than readthrough transcription from *CAV1*, we combined *de novo* transcript assembly with publicly available CAGE-seq (Figure 5a). Transcript assembly recovered the canonical *CAV1* mRNA and the novel downstream transcript, *eCAV1_WT*. The overlap of *eCAV1_WT* with a distinct CAGE-seq peak downstream of the assembled *CAV1* mRNA supports transcription initiation from a *de novo* TSS in the *CAV1* 3^*′*^ UTR. Together, these data substantiate the presence of bona fide enhancer activity at the *CAV1* 3^*′*^ UTR that drives eRNA transcription.

Having established enhancer activity at this locus, we investigated the potential impact of ALS-associated SNPs within the proximal *CAV1/CAV2* enhancer. By combining *in silico* predictions with experimental validation, we modelled how these enhancer variants may influence molecular function on multiple levels, including the impact on enhancer regulatory DNA and the eRNA transcribed from this locus. Even within the same enhancer element, ALS-associated SNPs may exert their effects through distinct pathways. This complexity is further compounded by the dual genic-regulatory context of 3^*′*^ UTR-associated enhancers. As such, we cannot conclude whether the predicted or observed impacts of ALS-associated SNPs reflect disruption of enhancer activity, 3^*′*^ UTR function, or both. However, explicitly disentangling enhancer from 3^*′*^ UTR function at this locus risks overlooking the integrated regulatory impact of this locus. The presence of five independent ALS-associated SNPs within the proximal *CAV1/CAV2* enhancer suggests that disruption of this region is functionally important, with individual variants potentially acting through distinct mechanisms - differentially impacting enhancer activity, 3^*′*^ UTR regulation, or both - yet converging on pathways relevant to ALS pathogenesis.

Screening ALS-associated SNPs within the proximal *CAV1/CAV2* enhancer via luciferase reporter assays demonstrated that enhancer variants can reduce enhancer regulatory activity in this system, with chr7:116,200,589T*>*A and chr7:116,200,705C*>*T significantly impairing the ability of the enhancer to drive reporter gene expression compared to the WT enhancer sequence. In contrast, our *in silico* predictions did not implicate these SNPs as potential disruptors of TF binding and eRNA structure, instead highlighting chr7:116,199,522T*>*C as a variant with potential multifactorial effects, predicted to impact enhancer function at both the DNA and RNA level.

To explore the RNA-level consequences of ALS-associated SNPs in the proximal *CAV1/CAV2* enhancer, we modelled the structure and stability of the antisense eRNA *eCAV1_WT*, assembled from SH-SY5Y TT-seq data, compared to *eCAV1_ALS*, generated by the *in silico* introduction of the chr7:116,199,522T*>*C substitution (Figure 5). The ALS variant induced predicted structural rearrangements and consistently increased predicted stability across multiple computational metrics, including more negative MFE and MFEden values and lower thermodynamic z-score minima. Structural modelling revealed the emergence of a hairpin loop structure in *eCAV1_ALS* absent from the WT structure, highlighting the ability of a single nucleotide substitution to reorganise local RNA folding and enhance overall folding stability. These observations provide a proof-of-concept that ALS-associated SNPs can substantially alter eRNA folding landscapes and underscore the importance of nucleotide order in shaping eRNA structure and stability.

Such structural rearrangements may influence eRNA half-life, RNA-protein interactions, and recruitment of chromatin regulators, potentially altering enhancer activity and downstream target gene expression. The expanded hairpin loop structure introduced by the ALS variant may have functional consequences, as such structural motifs can act as kinetic barriers to unfolding, enhancing overall RNA stability (Rissone et al., 2022), or provide platforms for structure-specific protein binding (reviewed in Svoboda and Di Cara, 2006; Harrison and Bose, 2022). For sequence-independent functions of eRNAs, such as CBP binding (Bose et al., 2017), the increased overall stability bestowed by the hairpin loop could enhance eRNA functionality by improving resistance to degradation. Thus disease-associated alterations in eRNA structure and stability may represent a mechanism by which non-coding SNPs contribute to disease risk. Experimental approaches such as road-block qPCR (Watson et al., 2020) or thiol(SH)-linked alkylation for the metabolic sequencing of RNA (SLAM-seq) (Herzog et al., 2017) could be employed to test whether ALS-associated SNPs indeed alter eRNA half-life, while future studies will be necessary to test whether these structural changes exist in a physiological context, impact eRNA function, enhancer activity, or target gene expression in neuronal contexts relevant to ALS. The proximal *CAV1/CAV2* enhancer may be more prone to DNA sequence-dependent changes in activity, due to the inherent evolutionary conservation of 3^*′*^ UTR sequences (Siepel et al., 2005; Xie et al., 2005). Disease-associated variants are enriched in evolutionarily conserved putative enhancer elements (Hujoel et al., 2019). Sequences under strong negative selection are more likely to harbour causal genetic variants, potentially due to the critical component of sequence in determining their function. Consequently, variants within conserved regions are more likely to exert a phenotypic effect, paralleling the impact observed for mutations in conserved protein-coding regions (Reva et al., 2011). Consistent with this, our analyses predict that the chr7:116,199,522T*>*C SNP in the proximal *CAV1/CAV2* enhancer may impair binding of MEF2 family transcription factors.

To model the potential impact of impaired MEF2D binding at this locus, we performed siRNA-mediated knockdown of *MEF2D* in SH-SY5Y cells, which resulted in a significant reduction in both *CAV1* mRNA and eRNA transcription from the proximal enhancer, highlighting the importance of MEF2 activity for robust transcription at this locus. Given that MEF2 can recruit epigenetic co-regulators, including CBP, to modulate chromatin structure (He et al., 2011; Youn et al., 2000), impaired MEF2 binding may reduce CBP recruitment, leading to altered chromatin accessibility, diminished RNAPII recruitment, and ultimately compromised enhancer activity. The similar reductions in *CAV1* mRNA and eRNA expression observed upon CBP/p300 inhibition (Figure 6b) and *MEF2D* knockdown (Figure 6d) support the notion that MEF2D and CBP/p300 may act in a coordinated manner to regulate activity at the proximal *CAV1/CAV2* enhancer. Future studies using genetically modified cell lines harbouring the chr7:116,199,522T*>*C SNP could directly test these predicted disruptions to TF binding and co-activator recruitment. Immunoprecipitation-based assays, such as ChIP-qPCR, would enable assessment of the impact of the SNP on MEF2 binding, CBP recruitment, and H3K27ac deposition at the proximal enhancer.

Collectively, our results highlight the proximal *CAV1/CAV2* enhancer as a functionally important regulatory element embedded within the 3^*′*^ UTR of the ALS risk gene, *CAV1*. This unique intragenic context provides a rare opportunity to link non-coding ALS-associated variants directly to both enhancer function and host gene regulation, offering mechanistic insight into how these SNPs may contribute to disease risk. In particular, the chr7:116,199,522T*>*C SNP exemplifies how a single variant can differentially impact multiple layers of regulation: altering enhancer DNA sequence to disrupt transcription factor binding, influencing recruitment of epigenetic co-regulators and chromatin remodelling, and reshaping the structure and stability of the transcribed eRNA. These multifactorial effects may converge to fine-tune *CAV1* expression, offering a mechanistic explanation for how non-coding variants within a single intragenic enhancer contribute to ALS pathogenesis. While the physiological impact of these changes remains to be established, our findings highlight the value of investigating non-coding regulatory elements - particularly intragenic enhancers within 3^*′*^ UTRs - as modulators of risk gene expression and potential contributors to ALS risk. Moreover, by focusing on a 3^*′*^ UTR-associated enhancer, we provide insight into an underexplored mechanism that may contribute to the unique transcriptional and regulatory profiles of 3^*′*^ UTRs, warranting future studies in disease-relevant models to assess their functional consequences *in vivo*.

## Materials and Methods

### Molecular biology and cloning

Polymerase chain reaction (PCR) was performed in a 50 *µ*l volume: 0.5 *µ*l PFuUltra II fusion HS DNA Polymerase (Agilent), 5% dimethyl sulfoxide (DMSO, Sigma), 200*µ*M dNTP mix, 0.5 *µ*M forward and reverse primer, and template DNA. DNA was denatured at 95 °C with an extension temperature of 72 °C, with 30 cycles. Successful assembly and base substitutions were confirmed by Sanger sequencing (Eurofins Genomics) with pre-mixed primers.

Wild-type and ALS-associated *CAV1* enhancer constructs were generated from a synthetic *CAV1* enhancer G-block (CAV1_enhancer_WT, Thermo Fisher GeneArt). ALS-associated mutations were introduced by site-directed mutagenesis, and fragments were cloned into the pGL3-Promoter vector (Promega, #E1761) using *Mlu*I and *Xho*I restriction sites. Plasmids used for CRISPRi were pHR-UCOE-SFFV-Zim3-dCas9-P2A-Hygro (Zim3-dCas9, Addgene #188768) (Replogle et al., 2022) and modified from lentiGuide-Puro (Addgene #52963) (Sanjana et al., 2014). sgRNAs were cloned into lentiGuide-Puro using a modified protocol from the Zhang lab (Sanjana et al., 2014; Shalem et al., 2014). Briefly, two oligonucleotides were designed for each sgRNA containing overhangs compatible with *BsmB*I digest of the lentiGuide-Puro vector. sgRNA sequences are listed in Table S1.

### Mammalian cell culture

SH-SY5Y cell cultures were maintained in Dulbecco’s Modified Eagle’s Medium (DMEM) (Cytiva) supplemented with 10% (v/v) heat-inactivated foetal bovine serum (FBS) (Thermo Fisher) and 50U/ml Penicillin/Streptomycin (Thermo Fisher) at 37 °C, 5% CO_2_. SH-SY5Y cell cultures stably expressing Zim3-dCas9 were maintained in DMEM supplemented with 10% (v/v) FBS and 500*µ*g/ml Hygromycin B (Thermo Fisher) at 37 °C, 5% CO_2_. HEK293FT cell cultures were maintained in DMEM supplemented with 10% (v/v) FBS and 50U/ml Penicillin/Streptomycin at 37 °C, 5% CO_2_.

### Transfection and transduction of plasmid DNA

Electroporation of SH-SY5Y cells with pGL3-Promoter (and derivatives, Promega #E1761), pGL3-Basic (Addgene #212936), pGL3-Control (Addgene #212937), and 759_Renilla (derived from pGL4.70[hRluc], Promega #E6881) was performed using the Neon^®^ Transfection System (Thermo Fisher) according to the manufacturer’s instructions. For luciferase reporter assays, transfections were performed in DMEM without phenol red (Cytiva, SH30284.01) supplemented with 10% FBS. For Dual-Glo luminescence experiments, vectors were co-transfected with 759_Renilla at a 1:1 ratio, with equal concentrations of plasmid used for each condition. For SH-SY5Ys the pulse conditions were set to 1200V, at a 20ms pulse width with 3 pulses. For generation of SH-SY5Y polyclonal lines stably expressing Zim3-dCas9 for CRISPRi and delivery of gRNA constructs (modified from lentiGuide-Puro), lentiviral packaging of transfer plasmids was performed in HEK293FT cells with the pMD2.G (VSVG) envelope (Addgene #12259) and psPAX2 (Addgene #12260) packaging plasmids. Transfections were carried out using LipoD293 (SignaGen Laboratories) according to the manufacturer’s instructions. Lentivirus was concentrated using Lenti-X concentrator (Takara Bio) and used to transduce SH-SY5Y cells in the presence of 8 *µ*g/ml polybrene.

### Antibiotic selection

For CRISPRi, SH-SY5Y cells expressing Zim3-dCas9 were selected with 500*µ*g/ml hygromycin. Following gRNA transduction, cells were selected with 500*µ*g/ml hygromycin and 1mg/ml puromycin for one week, followed by maintenance with 500*µ*g/ml hygromycin and 2mg/ml puromycin.

### Small molecule treatment

The small molecule A-485 (Stratech) was used to inhibit CBP/p300 histone acetyltransferase activity (Lasko et al., 2017). SH-SY5Y cells were treated with 5*µ*M A-485 for 48 hours, with PBS-treated cells as a negative control.

### siRNA treatment

siRNA-mediated knockdown was performed in SH-SY5Y cells using the Neon^®^ Transfection System (Thermo Fisher) as previously described. Cells were transfected with 150nM *MEF2D* siRNA (Silencer Select, Thermo Fisher), 150nM negative control siRNA (Silencer, Thermo Fisher), or PBS. RNA was extracted 48 hours post-transfection for RT-qPCR.

### Western blot

To validate stable expression of Zim3-dCas9 in transduced SH-SY5Y cells, a western blot probing for *α*-Cas9 was performed. SH-SY5Y cells were washed with PBS, harvested by centrifugation and the cell pellet was lysed in RIPA Lysis Buffer (50mM Tris HCl (Sigma) pH 8 at 4 °C, 100mM NaCl (Melford), 2mM MgCl_2_ (Thermo Fisher), 1% Triton X-100 (Sigma), 0.1% Sodium deoxycholate (Sigma), 0.1% Sodium Dodecyl Sulfate (SDS, Sigma)), supplemented with 1mM Dithiothreitol (DTT, Melford), 1X Halt phosphatase inhibitor cocktail (Thermo Fisher), 10mM sodium butyrate (Sigma) and 500U/100*µ*l Benzonase (Insight Biotech). 25-50*µ*g of protein lysate was prepared, with addition of 1X Invitrogen NuPAGE LDS Sample Buffer (Thermo Fisher), and loaded onto a NuPAGE 3-8% Tris-Acetate gel (Thermo Fisher), and run using 1X Tris-Acetate SDS running buffer, with 500*µ*l Invitrogen NuPAGE Antioxidant (Thermo Fisher). Gel was transferred onto nitrocellulose membrane using the semi-dry Trans-Blot Turbo Transfer System (Bio-Rad) and the Trans-Blot Turbo Mini 0.2*µ*m nitrocellulose transfer packs (Bio-Rad), before blocking and probing for *α*-GAPDH (1:10,000, Proteintech, 60004-1-Ig) and *α*-Cas9 (1:500, Abcam, ab191468). Western blots were imaged using G-BOX Chem-XRQ (Syngene) and GeneSys software. Images were processed with ImageJ.

### Luciferase reporter assays

24 hours following transfection of luciferase reporter constructs into SH-SY5Y cells, firefly and *Renilla* luciferase activity was measured using the Promega Dual-Glo luciferase assay system (Promega) according to the manufacturer’s instructions. Luminescence subsequently measured using Hidex Sense plate reader and luminescence settings (orbital shake with 5 second duration, IR cutoff filter and 5 second counting time). Relative luciferase activity was calculated as ratio of firefly to *Renilla* signal and normalised to PBS-treated controls. Error bars were calculated using standard error. Significance of results were determined by one-way ANOVA test using the `aov` function in `R`, followed by computation of Tukey Honest Significant Differences using the `TukeyHSD` function.

### Quantitative reverse transcription PCR (RT-qPCR)

Total RNA was extracted from SH-SY5Y cells using TRIzol reagent (Thermo Fisher) with chloroform phase separation and isopropanol precipitation. RNA concentration was quantified using Qubit RNA Broad Range assay kit (Thermo Fisher) and a Qubit 4 fluorometer (Thermo Fisher). 2*µ*g of RNA per sample was taken to generate cDNA by reverse transcription. Reverse transcription was performed using either the High-Capacity cDNA kit (Thermo Fisher) or the High-Capacity RNA-to-cDNA kit (Thermo Fisher), according to manufacturer’s guidelines. RT-qPCR was performed using Power SYBR Green PCR Master Mix (Thermo Fisher) on a QuantStudio 12K Flex system. Primers used for target genes, including *CAV1* coding sequences and eRNAs, are listed in Table S2. Primer concentrations ranged from 100–600 nM. Relative gene expression was calculated using the ΔΔCt method with 18S rRNA as an internal control. All experiments were performed at least in triplicate. Error bars represent standard error of the mean. Statistical significance was determined using unpaired moderated t-tests with the `limma eBayes` (Ritchie et al., 2015) function or paired t-tests using the `ttest` function in `R`.

### Published dataset sources

In addition to iPSC-derived motor neuron datasets, we also utilised published datasets from WT SK-N-SH and SH-SY5Y neuroblastoma cells. A summary of the sources of published datasets, including accessions is provided in Table 1.

### Hi-C

*In situ* Hi-C contact matrices for *in vitro* differentiated motor neurons (Zhang et al., 2022) were downloaded from the ENCODE portal; accession numbers are listed in Table 1 (Luo et al., 2020; Hitz et al., 2023; ENCODE Project Consortium, 2012). The `HiCExplorer` command line suite (Ramírez et al., 2018) was utilised to convert file formats using the tool `hicConvertFormat`. Contact domains and TAD boundaries were identified using the tool `hicFindTADs` and the flag `–correctForMultipleTesting fdr`. For downstream analysis of chromosome topology in ALS patients, Hi-C contact matrices from ALS patient derived motor neurons were combined using the `HiCExplorer` (Ramírez et al., 2018) tool `hicSumMatrices`. Configuration files for plotting Hi-C track lines were generated using the `make_tracks_file` function from the `pyGenomeTracks` package. Subsequently track lines were plotted using `hicPlotTADs` tool from `HiCExplorer` (Ramírez et al., 2018) with the hg38 plotting region defined using the `–region` flag. To generate heatmaps demonstrating CTCF binding at TAD boundaries, CTCF ChIP-seq data from SK-N-SH neuroblastoma cells was downloaded from the ENCODE portal with accession number listed in Table 1 (Luo et al., 2020; Hitz et al., 2023; ENCODE Project Consortium, 2012). Matrices of CTCF signal around TAD boundaries were computed using the `deepTools` suite with the tool `computeMatrix` with the `scale-regions` parameter and the flags `-a 500 -b 500`. Heatmaps were plotted using the `deepTools` tool `plotHeatmap` (Ramírez et al., 2016).

### Assay for transposase-accessible chromatin with sequencing (ATAC-seq)

ATAC-seq datasets from iPSC-derived motor neurons from healthy controls and ALS patients (Zhang et al., 2022) were downloaded from the ENCODE portal; accession numbers are listed in Table 1 (Luo et al., 2020; Hitz et al., 2023; ENCODE Project Consortium, 2012).

### Chromatin immunoprecipitation sequencing (ChIP-seq)

H3K27ac and H3K4me1 datasets, including pseudoreplicated peak and fold change over control files, from iPSC-derived motor neurons from healthy controls and ALS patients (Zhang et al., 2022) were downloaded from the ENCODE portal; accession numbers are listed in Table 1 (Luo et al., 2020; Hitz et al., 2023; ENCODE Project Consortium, 2012).

### RNA-sequencing (RNA-seq)

Total SH-SY5Y RNA-seq raw fastq files (Liu et al., 2022) were downloaded from the NCBI GEO database, with accession number listed in Table 1 (Edgar et al., 2002), and aligned to the hg19 reference genome using the `STAR` aligner (Dobin et al., 2013) with the flags `–outSAMtype BAM SortedByCoordinate –quantMode TranscriptomeSAM –outFilterType BySJout –peOverlapNbasesMin` `40 –peOverlapMMp 0.8`.

### Cap analysis of gene expression sequencing (CAGE-seq)

Cap analysis of gene expression sequencing (CAGE-seq) dataset from SK-N-SH cells was downloaded from the ENCODE portal with accession number listed in Table 1 (Luo et al., 2020; Hitz et al., 2023; ENCODE Project Consortium, 2012).

### Visualisation of next-generation sequencing data

For visualisation of output bigWig files and transcript GTF files, tracklines were generated using `pyGenomeTracks` (Lopez-Delisle et al., 2021; Ramírez et al., 2018), with gene locations provided by `TxDb.Hsapiens.UCSC.hg19.knownGene` (Carlson and Maintainer, 2015).

### Cleavage under targets and release using nuclease (CUT&RUN) sequencing

Cleavage under targets and release using nuclease (CUT&RUN) sequencing libraries were generated using the CUT&RUN kit from Active Motif, as per manufacturer’s guidelines. Isolated nuclei from 500,000 SH-SY5Y cells were utilised as input, using 1*µ*g *α*-Histone H3 (Abcam, ab1791) as the control antibody and 1*µ*g *α*-Histone H3 acetyl K27 (Abcam, ab4729) as the target antibody. Library preparation was performed using the NEBNext® Ultra™ II kit. Following library preparation, an additional single-sided cleanup was performed to remove sub-nucleosomal peaks. Prior to sequencing, library preparation size distribution was examined using TapeStation 4150 system (Agilent) and High Sensitivity DNA ScreenTape (Agilent). Paired-end (PE150) sequencing of libraries was performed by Novogene using Illumina V1.5 reagents and the Illumina NovaSeq6000 platform, generating 9GB raw data. Quality control and adapter trimming of raw fastq files was performed using `Trim Galore`(Krueger, 2012). Reads were aligned to the hg19 reference genome using `bowtie2` (Langmead and Salzberg, 2012) using the flags `–local –very-sensitive-local` `-I 10 -X 700 –dovetail`. Aligned reads were filtered using `samtools view`, and sorted using `samtools sort` (Li et al., 2009). CUT&RUN peaks were called from the alignment BAM files using `MACS3 callpeak` (Zhang et al., 2008) using histone H3 as the control file and H3K27ac as the treatment file. Subsequently H3K27ac fold enrichment over H3 control was calculated using `MACS3 bdgcmp` (Zhang et al., 2008) using the flag `-m FE`, to allow for comparison to published ChIP-seq datasets.

### Transient transcriptome sequencing (TT-seq)

TT-seq libraries were generated using a protocol modified from Gregersen et al. (2020). Briefly, SH-SY5Y cells were labelled with 500*µ*M 4-thiouridine (4sU, Scientific Laboratory Supplies) for 5 minutes at 37 °C, 5% CO_2_. Cells were lysed in TRIzol (Thermo Fisher) and total RNA was extracted using chloroform phase separation and isopropanol precipitation. For normalisation purposes, 500ng of *S. cerevisiae* 4-thiouracil (4TU, Sigma)-labelled RNA was added to 100*µ*g of 4sU-labelled RNA. RNA was chemically fragmented using 20*µ*l of 1M NaOH (Thermo Fisher), and 4sU-labelled RNA was biotinylated with MTSEA biotin-XX (Biotium) and purified. Incorporation for 4sU was assessed by RNA dot blot, probed with streptavidin-HRP (1:1,000, Epigentek Group Inc). Biotinylated RNA was enriched by streptavidin pull-down using *µ*MACS Streptavidin MicroBeads (Miltenyi Biotec) and eluted in RNase-free water. The size distribution of 4sU-RNA was examined using TapeStation 4150 system (Agilent) and High Sensitivity RNA ScreenTape (Agilent), and RNA concentration was measured using Qubit RNA HS Assay kit (Thermo Fisher) and a Qubit 4 fluorometer (Thermo Fisher). Library preparation was performed using Lexogen CORALL Total RNA-seq Library Prep Kit (V1 and V2 with UDI 2nt Set A1) with UDIs (Lexogen) as per manufacturer’s guidelines. Sequencing library size distribution was examined using TapeStation 4150 system (Agilent) and High Sensitivity DNA ScreenTape (Agilent). Paired-end (PE150) sequencing of libraries was performed by Novogene using Illumina V1.5 reagents and the Illumina NovaSeq6000 platform, generating 30GB of raw data. Quality control on raw fastq files was performed using `fastqc`(Andrews, 2010). Unique molecular identifier (UMI) sequences were extracted and appended to header using `UMI-tools extract`(Smith et al., 2017). Adapters were trimmed and post-trimming quality control was performed using `Trim Galore`(Krueger, 2012). Trimmed reads were aligned to the hg19 human reference genome and the sacCer3 yeast reference genome using `STAR` aligner (Dobin et al., 2013) using the flags `–outSAMtype BAM SortedByCoordinate` `–quantMode GeneCounts –outFilterType BySJout` `–peOverlapNbasesMin 40 –peOverlapMMp 0.8`. Post-alignment quality control, BAM file sorting, and BAM file indexing was performed using `samtools` (Li et al., 2009). Reads were grouped based on their UMI and mapping coordinates using `UMI-tools group` and then deduplicated using `UMI-tools dedup` (Smith et al., 2017). Normalisation was performed by calculating scaling factors based on exogenous yeast spike-in read count. Scaling factors were calculated using `DESeq2` (Love et al., 2014) and the function `estimateSizeFactors` to generate size factors using `ReadsPerGene.out.tab` output files from the sacCer3 alignment generated by `STAR` aligner. To identify correlation between replicates, a `DESeq2` dataset was generated using the `ReadsPerGene.out.tab` output files from the hg19 alignment generated by `STAR` aligner. The `DESeq2` dataset was normalised using spike-in scaling factors using the `sizeFactors` function, and normalised gene counts were generated using the `counts` function with the argument `normalized = TRUE`. Correlation matrices were then calculated using the base `R` function `cor` with the argument `method = “pearson”`. Stranded reads files were generated using the `samtools` function `view` (Li et al., 2009). Stranded BAM files were scaled based on exogenous spike-in using the `bamCoverage` function from `deepTools` (Ramírez et al., 2016) using the `–scaleFactor` flag and scaling factors calculated using `DESeq2`(Love et al., 2014). To assemble RNA transcripts from TT-seq data, *de novo* transcript assembly was performed using `StringTie` with the flags `–fr -m 50 -s 1 -c 1` `-j 1` (Pertea et al., 2015). Novel eRNA transcripts were then identified using the package `gffcompare` (Pertea and Pertea, 2020), comparing the output transcript assembly GTF file to the reference hg19 GENCODE annotation GTF (Frankish et al., 2019). For downstream analyses, transcripts that were transcribed from within, and restricted to candidate *CAV1/CAV2* enhancer regions with the class codes ‘u’, ‘e’, ‘x’ and ‘o’ were considered to be potential eRNAs.

### RNA structure prediction

MFE eRNA structures were predicted using `RNAfold` with the flags `-p –noLP –salt 0.1` (Lorenz et al., 2011), and visualised using forna from ViennaRNA (Lorenz et al., 2011). Thermodynamic z-scores of RNA sequences were calculated using `ScanFold` (Andrews et al., 2018), with a window size of 10nt and a step size of 1nt for 100 randomisations.. To calculate the MFEden as function of GC-content, shuffled RNA sequences were generated with lengths ranging from 50-1000 nt (50 nt increments) and GC-contents of 20%, 40%, 50%, 60%, 70%, and 80%, with 100 sequences per condition. MFE for each sequence was calculated using `RNAfold`. For each sequence length, MFEref_*L*_ was computed by generating shuffled RNA sequences with equimolar nucleotide composition, which were then processed with `RNAfold`. MFEden values for the shuffled sequences were calculated using a custom `bash` script. A second-degree polynomial regression model of MFEden versus GC-content was fitted in `R` using the `lm` function with a quadratic term. The model was trained on the shuffled sequence dataset and evaluated on an independent test set. Model residuals were calculated in `R` using the `residuals` function and examined for correlation with GC content to assess potential bias. Shuffled RNA sequences with the same nucleotide composition as the native sequence were generated using the bash command `shuf`, and the output sequences were processed with `RNAfold` for MFE calculation.

### Per gene exonic versus intronic read coverage

To compare coverage over intronic versus exonic regions for TT-seq and total RNA-seq, intronic features were extracted from the GENCODE v19 annotation using the `R` package `GenomicFeatures` (Lawrence et al., 2013). We extracted the disjoint intronic parts using `intronicParts` and the resultant object was subsetted to remove any spurious introns larger than 5kb. Read counts over exons were obtained using `featureCounts` with the flags `-p -F GTF -t exon` with the GENCODE V19 as the input annotation file. Read counts over introns were obtained using `featureCounts` with the flags `-p -F GTF -t sequence_feature` and the intron annotation file generated using `GenomicFeatures` as the input annotation file. The ratio of exonic to intronic reads were then calculated and visualised using `ggplot2` (Wickham, 2016).

### Transcription factor binding motifs

Transcription factor binding motifs within *CAV1/CAV2* candidate enhancers were identified using the JASPAR 2024 Transcription Factor Binding Site database (JASPAR CORE 2024) (Rauluseviciute et al., 2024) trackline on the UCSC genome browser (Raney et al., 2024).

## Supporting information

Fig. S2: Residuals of the polynomial regression for MFEden versus GC-content.

Fig. S1: CTCF binds at TAD boundaries. CTCF binding as determined by ChIP-seq signal in SK-N-SH cells across all TADs genome-wide.

## Acknowledgments

D.A.B is funded by the Wellcome Trust and the Royal Society (grant numbers 213501/Z/18/Z and 220192/Z/20/Z respectively); L.J.H was supported by a studentship from the Medical Research Council Discovery Medicine North (DiMeN) Doctoral Training Partnership (grant number: MR/N013840); J.C-K was supported by the Wellcome Trust (grant number: 216596/Z/19/Z), the ALS Association (23-PP-664), and TargetALS; D.A.B also received funding from The Royal Society (grant number: RSG\R1\180410) and a BBSRC Pioneer Award (grant number: BB/Y513453/1). This research was funded in whole, or in part, by the Wellcome Trust [213501/Z/18/Z, 220192/Z/20/Z and 216596/Z/19/Z]. For the purpose of Open Access, the author has applied a CC BY public copyright licence to any Author Accepted Manuscript version arising from this submission.

## Author contributions

The study was conceived and led by D.A.B, L.J.H and J.C-K. D.A.B and L.J.H designed and carried out all experiments, including analysis of NGS data, with input from J.C-K and T.M. L.J.H and D.A.B. wrote the manuscript; All authors reviewed and commented on the manuscript.

## Supplementary materials

**Fig. S1:**
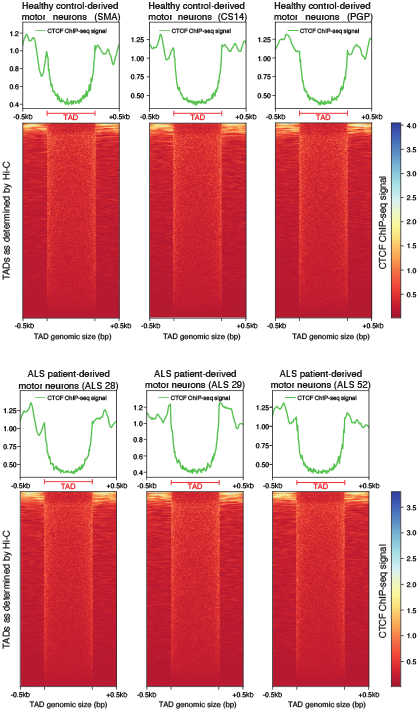
CTCF binds at TAD boundaries. CTCF binding as determined by ChIP-seq signal in SK-N-SH cells across all TADs genome-wide. TAD boundaries are determined by Hi-C contact frequencies in fibroblast-derived iPSCs differentiated *in vitro* into motor neurons (Luo et al., 2020; Hitz et al., 2023; ENCODE Project Consortium, 2012; Zhang et al., 2022).

**Fig. S2:**
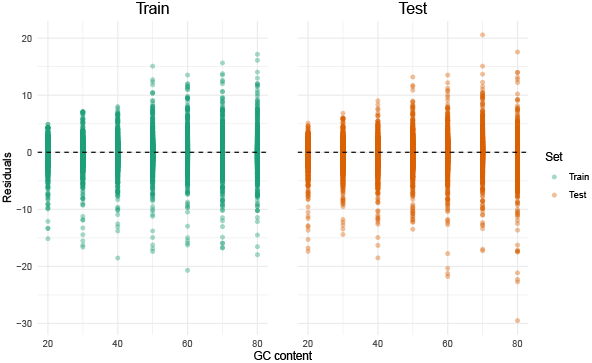
Residuals of the polynomial regression for MFEden versus GC-content. Residuals from the polynomial regression model are shown for the training (left) and independent test set (right). Residuals are centered near zero and display negligible correlation with GC content, indicating minimal bias.

**Table S1.**
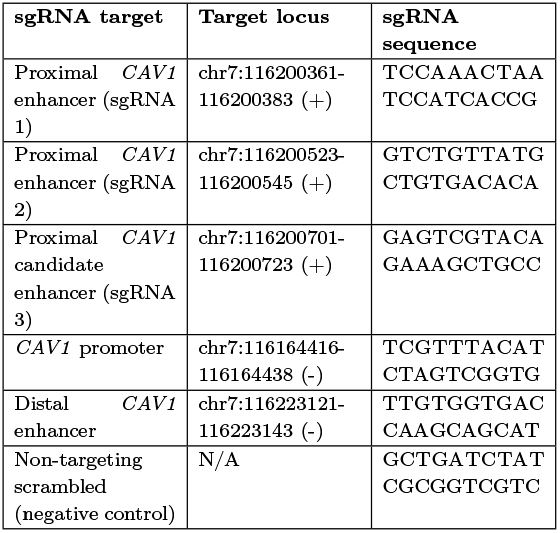
sgRNA targets and sequences used for CRISPRi in SH-SY5Y cells.

**Table S2.**
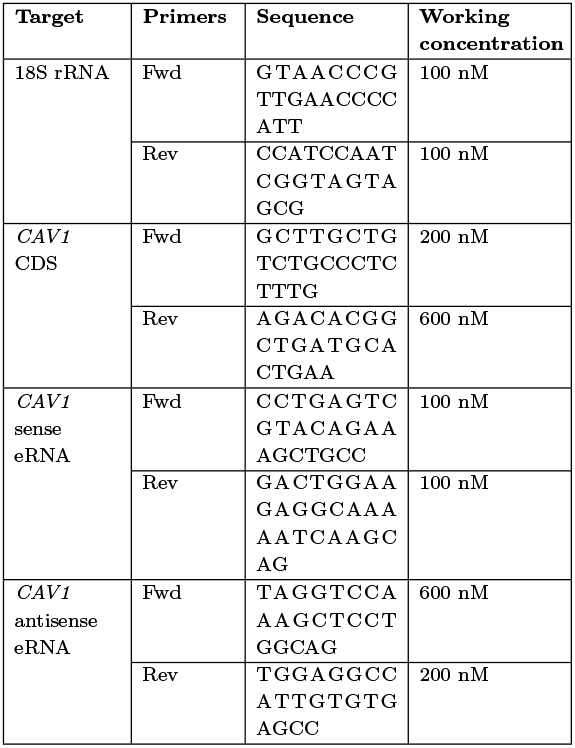
Primers used for RT-qPCR.

## References

Adey, B. N., Cooper-Knock, J., Al Khleifat, A., Fogh, I., van Damme, P., Corcia, P., Couratier, P., Hardiman, O., McLaughlin, R., Gotkine, M., Drory, V., Silani, V., Ticozzi, N., Veldink, J. H., van den Berg, L. H., de Carvalho, M., Pinto, S., Mora Pardina, J. S., Povedano Panades, M., Andersen, P. M., Weber, M., Başak, N. A., Shaw, C. E., Shaw, P. J., Morrison, K. E., Landers, J. E., Glass, J. D., Vourc’h, P., Dobson, R. J. B., Breen, G., Al-Chalabi, A., Jones, A. R., and Iacoangeli, A. (2023). Large-scale analyses of CAV1 and CAV2 suggest their expression is higher in post-mortem ALS brain tissue and affects survival. Front. Cell. Neurosci., 17:1112405.

Andrews, R. J., Roche, J., and Moss, W. N. (2018). ScanFold: an approach for genome-wide discovery of local RNA structural elements-applications to zika virus and HIV. PeerJ, 6(e6136):e6136.

Andrews, R. J., Rouse, W. B., O’Leary, C. A., Booher, N. J., and Moss, W. N. (2022). ScanFold 2.0: a rapid approach for identifying potential structured RNA targets in genomes and transcriptomes. PeerJ, 10:e14361.

Andrews, S. (2010). Fastqc: A quality control tool for high throughput sequence data. Accessed: 2025-09-08.

Babak, T., Blencowe, B. J., and Hughes, T. R. (2007). Considerations in the identification of functional RNA structural elements in genomic alignments. BMC Bioinformatics, 8(1):33.

Bal, E., Kumar, R., Hadigol, M., Holmes, A. B., Hilton, L. K., Loh, J. W., Dreval, K., Wong, J. C. H., Vlasevska, S., Corinaldesi, C., Soni, R. K., Basso, K., Morin, R. D., Khiabanian, H., Pasqualucci, L., and Dalla-Favera, R. (2022). Super-enhancer hypermutation alters oncogene expression in B cell lymphoma. Nature, pages 1–8.

Bonnet, E., Wuyts, J., Rouzé, P., and Van de Peer, Y. (2004). Evidence that microRNA precursors, unlike other non-coding RNAs, have lower folding free energies than random sequences. Bioinformatics, 20(17):2911–2917.

Bose, D. A., Donahue, G., Reinberg, D., Shiekhattar, R., Bonasio, R., and Berger, S. L. (2017). RNA binding to CBP stimulates histone acetylation and transcription. Cell, 168(1-2):135–149.e22.

Bäumer, D., Talbot, K., and Turner, M. R. (2014). Advances in motor neurone disease. J. R. Soc. Med., 107(1):14–21.

Carlson, M. and Maintainer, B. P. (2015). TxDb. hsapiens. UCSC. hg19. knowngene. Bioconductor.

Carullo, N. V. N. and Day, J. J. (2019). Genomic enhancers in brain health and disease. Genes, 10(1).

Clote, P., Ferré, F., Kranakis, E., and Krizanc, D. (2005). Structural RNA has lower folding energy than random RNA of the same dinucleotide frequency. RNA, 11(5):578–591.

Cooper-Knock, J., Zhang, S., Kenna, K. P., Moll, T., Franklin, J. P., Allen, S., Nezhad, H. G., Iacoangeli, A., Yacovzada, N. Y., Eitan, C., Hornstein, E., Elhaik, E., Celadova, P., Bose, D., Farhan, S., Fishilevich, S., Lancet, D., Morrison, K. E., Shaw, C. E., Al-Chalabi, A., Project MinE ALS Sequencing Consortium, Veldink, J. H., Kirby, J., Snyder, M. P., and Shaw, P. J. (2021). Rare variant burden analysis within enhancers identifies CAV1 as an ALS risk gene. Cell Rep., 34(5):108730.

Creyghton, M. P., Cheng, A. W., Welstead, G. G., Kooistra, T., Carey, B. W., Steine, E. J., Hanna, J., Lodato, M. A., Frampton, G. M., Sharp, P. A., Boyer, L. A., Young, R. A., and Jaenisch, R. (2010). Histone H3K27ac separates active from poised enhancers and predicts developmental state. Proc. Natl. Acad. Sci. U. S. A., 107(50):21931–21936.

De Santa, F., Barozzi, I., Mietton, F., Ghisletti, S., Polletti, S., Tusi, B. K., Muller, H., Ragoussis, J., Wei, C.-L., and Natoli, G. (2010). A large fraction of extragenic RNA pol II transcription sites overlap enhancers. PLoS Biol., 8(5):e1000384.

Dixon, J. R., Jung, I., Selvaraj, S., Shen, Y., Antosiewicz-Bourget, J. E., Lee, A. Y., Ye, Z., Kim, A., Rajagopal, N., Xie, W., Diao, Y., Liang, J., Zhao, H., Lobanenkov, V. V., Ecker, J. R., Thomson, J. A., and Ren, B. (2015). Chromatin architecture reorganization during stem cell differentiation. Nature, 518(7539):331–336.

Dixon, J. R., Selvaraj, S., Yue, F., Kim, A., Li, Y., Shen, Y., Hu, M., Liu, J. S., and Ren, B. (2012). Topological domains in mammalian genomes identified by analysis of chromatin interactions. Nature, 485(7398):376–380.

Dobin, A., Davis, C. A., Schlesinger, F., Drenkow, J., Zaleski, C., Jha, S., Batut, P., Chaisson, M., and Gingeras, T. R. (2013). STAR: ultrafast universal RNA-seq aligner. Bioinformatics, 29(1):15–21.

Dowen, J. M., Fan, Z. P., Hnisz, D., Ren, G., Abraham, B. J., Zhang, L. N., Weintraub, A. S., Schujiers, J., Lee, T. I., Zhao, K., and Young, R. A. (2014). Control of cell identity genes occurs in insulated neighborhoods in mammalian chromosomes. Cell, 159(2):374–387.

Edgar, R., Domrachev, M., and Lash, A. E. (2002). Gene expression omnibus: NCBI gene expression and hybridization array data repository. Nucleic Acids Res., 30(1):207–210.

Egawa, J., Zemljic-Harpf, A., Mandyam, C. D., Niesman, I. R., Lysenko, L. V., Kleschevnikov, A. M., Roth, D. M., Patel, H. H., Patel, P. M., and Head, B. P. (2018). Neuron-targeted caveolin-1 promotes ultrastructural and functional hippocampal synaptic plasticity. Cereb. Cortex, 28(9):3255–3266.

ENCODE Project Consortium (2012). An integrated encyclopedia of DNA elements in the human genome. Nature, 489(7414):57– 74.

Ernst, J., Kheradpour, P., Mikkelsen, T. S., Shoresh, N., Ward, L. D., Epstein, C. B., Zhang, X., Wang, L., Issner, R., Coyne, M., Ku, M., Durham, T., Kellis, M., and Bernstein, B. E. (2011). Mapping and analysis of chromatin state dynamics in nine human cell types. Nature, 473(7345):43–49.

Farh, K. K.-H., Marson, A., Zhu, J., Kleinewietfeld, M., Housley, W. J., Beik, S., Shoresh, N., Whitton, H., Ryan, R. J. H., Shishkin, A. A., Hatan, M., Carrasco-Alfonso, M. J., Mayer, D., Luckey, C. J., Patsopoulos, N. A., De Jager, P. L., Kuchroo, V. K., Epstein, C. B., Daly, M. J., Hafler, D. A., and Bernstein, B. E. (2015). Genetic and epigenetic fine mapping of causal autoimmune disease variants. Nature, 518(7539):337–343.

Finucane, H. K., Bulik-Sullivan, B., Gusev, A., Trynka, G., Reshef, Y., Loh, P.-R., Anttila, V., Xu, H., Zang, C., Farh, K., Ripke, S., Day, F. R., ReproGen Consortium, Schizophrenia Working Group of the Psychiatric Genomics Consortium, RACI Consortium, Purcell, S., Stahl, E., Lindstrom, S., Perry, J. R. B., Okada, Y., Raychaudhuri, S., Daly, M. J., Patterson, N., Neale, B. M., and Price, A. L. (2015). Partitioning heritability by functional annotation using genome-wide association summary statistics. Nat. Genet., 47(11):1228– 1235.

Flavell, S. W., Cowan, C. W., Kim, T.-K., Greer, P. L., Lin, Y., Paradis, S., Griffith, E. C., Hu, L. S., Chen, C., and Greenberg, M. E. (2006). Activity-dependent regulation of MEF2 transcription factors suppresses excitatory synapse number. Science, 311(5763):1008–1012.

Frankish, A., Diekhans, M., Ferreira, A.-M., Johnson, R., Jungreis, I., Loveland, J., Mudge, J. M., Sisu, C., Wright, J., Armstrong, J., Barnes, I., Berry, A., Bignell, A., Carbonell Sala, S., Chrast, J., Cunningham, F., Di Domenico, T., Donaldson, S., Fiddes, I. T., García Girón, C., Gonzalez, J. M., Grego, T., Hardy, M., Hourlier, T., Hunt, T., Izuogu, O. G., Lagarde, J., Martin, F. J., Martínez, L., Mohanan, S., Muir, P., Navarro, F. C. P., Parker, A., Pei, B., Pozo, F., Ruffier, M., Schmitt, B. M., Stapleton, E., Suner, M.-M., Sycheva, I., Uszczynska-Ratajczak, B., Xu, J., Yates, A., Zerbino, D., Zhang, Y., Aken, B., Choudhary, J. S., Gerstein, M., Guigó, R., Hubbard, T. J. P., Kellis, M., Paten, B., Reymond, A., Tress, M. L., and Flicek, P. (2019). GENCODE reference annotation for the human and mouse genomes. Nucleic Acids Res., 47(D1):D766–D773.

Fulco, C. P., Nasser, J., Jones, T. R., Munson, G., Bergman, D. T., Subramanian, V., Grossman, S. R., Anyoha, R., Doughty, B. R., Patwardhan, T. A., Nguyen, T. H., Kane, M., Perez, E. M., Durand, N. C., Lareau, C. A., Stamenova, E. K., Aiden, E. L., Lander, E. S., and Engreitz, J. M. (2019). Activity-by-contact model of enhancer–promoter regulation from thousands of CRISPR perturbations. Nat. Genet., 51(12):1664–1669.

Gong, X., Tang, X., Wiedmann, M., Wang, X., Peng, J., Zheng, D., Blair, L. A. C., Marshall, J., and Mao, Z. (2003). Cdk5-mediated inhibition of the protective effects of transcription factor MEF2 in neurotoxicity-induced apoptosis. Neuron, 38(1):33–46.

Gossett, L. A., Kelvin, D. J., Sternberg, E. A., and Olson, E. N. (1989). A new myocyte-specific enhancer-binding factor that recognizes a conserved element associated with multiple muscle-specific genes. Mol. Cell. Biol., 9(11):5022–5033.

Gregersen, L. H., Mitter, R., and Svejstrup, J. Q. (2020). Using TTchem-seq for profiling nascent transcription and measuring transcript elongation. Nat. Protoc., 15(2):604–627.

Hah, N., Danko, C. G., Core, L., Waterfall, J. J., Siepel, A., Lis, J. T., and Kraus, W. L. (2011). A rapid, extensive, and transient transcriptional response to estrogen signaling in breast cancer cells. Cell, 145(4):622–634.

Harrison, L. J. and Bose, D. (2022). Enhancer RNAs step forward: new insights into enhancer function. Development, 149(16).

He, J., Ye, J., Cai, Y., Riquelme, C., Liu, J. O., Liu, X., Han, A., and Chen, L. (2011). Structure of p300 bound to MEF2 on DNA reveals a mechanism of enhanceosome assembly. Nucleic Acids Res., 39(10):4464–4474.

Head, B. P., Hu, Y., Finley, J. C., Saldana, M. D., Bonds, J. A., Miyanohara, A., Niesman, I. R., Ali, S. S., Murray, F., Insel, P. A., Roth, D. M., Patel, H. H., and Patel, P. M. (2011). Neuron-targeted caveolin-1 protein enhances signaling and promotes arborization of primary neurons. J. Biol. Chem., 286(38):33310–33321.

Head, B. P., Patel, H. H., Tsutsumi, Y. M., Hu, Y., Mejia, T., Mora, R. C., Insel, P. A., Roth, D. M., Drummond, J. C., and Patel, P. M. (2008). Caveolin-1 expression is essential for N-methyl-D-aspartate receptor-mediated src and extracellular signal-regulated kinase 1/2 activation and protection of primary neurons from ischemic cell death. FASEB J., 22(3):828–840.

Head, B. P., Peart, J. N., Panneerselvam, M., Yokoyama, T., Pearn, M. L., Niesman, I. R., Bonds, J. A., Schilling, J. M., Miyanohara, A., Headrick, J., Ali, S. S., Roth, D. M., Patel, P. M., and Patel, H. H. (2010). Loss of caveolin-1 accelerates neurodegeneration and aging. PLoS One, 5(12):e15697.

Heintzman, N. D., Stuart, R. K., Hon, G., Fu, Y., Ching, C. W., Hawkins, R. D., Barrera, L. O., Van Calcar, S., Qu, C., Ching, K. A., Wang, W., Weng, Z., Green, R. D., Crawford, G. E., and Ren, B. (2007). Distinct and predictive chromatin signatures of transcriptional promoters and enhancers in the human genome. Nat. Genet., 39(3):311–318.

Herzog, V. A., Reichholf, B., Neumann, T., Rescheneder, P., Bhat, P., Burkard, T. R., Wlotzka, W., von Haeseler, A., Zuber, J., and Ameres, S. L. (2017). Thiol-linked alkylation of RNA to assess expression dynamics. Nat. Methods, 14(12):1198–1204.

Hilton, I. B., D’Ippolito, A. M., Vockley, C. M., Thakore, P. I., Crawford, G. E., Reddy, T. E., and Gersbach, C. A. (2015). Epigenome editing by a CRISPR-Cas9-based acetyltransferase activates genes from promoters and enhancers. Nat. Biotechnol., 33(5):510–517.

Hindorff, L. A., Sethupathy, P., Junkins, H. A., Ramos, E. M., Mehta, J. P., Collins, F. S., and Manolio, T. A. (2009). Potential etiologic and functional implications of genome-wide association loci for human diseases and traits. Proc. Natl. Acad. Sci. U. S. A., 106(23):9362–9367.

Hitz, B. C., Lee, J.-W., Jolanki, O., Kagda, M. S., Graham, K., Sud, P., Gabdank, I., Seth Strattan, J., Sloan, C. A., Dreszer, T., Rowe, L. D., Podduturi, N. R., Malladi, V. S., Chan, E. T., Davidson, J. M., Ho, M., Miyasato, S., Simison, M., Tanaka, F., Luo, Y., Whaling, I., Hong, E. L., Lee, B. T., Sandstrom, R., Rynes, E., Nelson, J., Nishida, A., Ingersoll, A., Buckley, M., Frerker, M., Kim, D. S., Boley, N., Trout, D., Dobin, A., Rahmanian, S., Wyman, D., Balderrama-Gutierrez, G., Reese, F., Durand, N. C., Dudchenko, O., Weisz, D., Rao, S. S. P., Blackburn, A., Gkountaroulis, D., Sadr, M., Olshansky, M., Eliaz, Y., Nguyen, D., Bochkov, I., Shamim, M. S., Mahajan, R., Aiden, E., Gingeras, T., Heath, S., Hirst, M., James Kent, W., Kundaje, A., Mortazavi, A., Wold, B., and Michael Cherry, J. (2023). The ENCODE uniform analysis pipelines.

Hofacker, I. L., Fontana, W., Stadler, P. F., Bonhoeffer, L. S., Tacker, M., and Schuster, P. (1994). Fast folding and comparison of RNA secondary structures. Monatshefte für Chemie /Chemical Monthly, 125(2):167–188.

Hujoel, M. L. A., Gazal, S., Hormozdiari, F., van de Geijn, B., and Price, A. L. (2019). Disease heritability enrichment of regulatory elements is concentrated in elements with ancient sequence age and conserved function across species. Am. J. Hum. Genet., 104(4):611–624.

Ji, X., Dadon, D. B., Powell, B. E., Fan, Z. P., Borges-Rivera, D., Shachar, S., Weintraub, A. S., Hnisz, D., Pegoraro, G., Lee, T. I., Misteli, T., Jaenisch, R., and Young, R. A. (2016). 3D chromosome regulatory landscape of human pluripotent cells. Cell Stem Cell, 18(2):262–275.

Kaikkonen, M. U., Spann, N. J., Heinz, S., Romanoski, C. E., Allison, K. A., Stender, J. D., Chun, H. B., Tough, D. F., Prinjha, R. K., Benner, C., and Glass, C. K. (2013). Remodeling of the enhancer landscape during macrophage activation is coupled to enhancer transcription. Mol. Cell, 51(3):310–325.

Kim, T.-K., Hemberg, M., Gray, J. M., Costa, A. M., Bear, D. M., Wu, J., Harmin, D. A., Laptewicz, M., Barbara-Haley, K., Kuersten, S., Markenscoff-Papadimitriou, E., Kuhl, D., Bito, H., Worley, P. F., Kreiman, G., and Greenberg, M. E. (2010). Widespread transcription at neuronal activity-regulated enhancers. Nature, 465(7295):182–187.

Kristjánsdóttir, K., Dziubek, A., Kang, H. M., and Kwak, H. (2020). Population-scale study of eRNA transcription reveals bipartite functional enhancer architecture. Nat. Commun., 11(1):5963.

Krueger, F. (2012). Trim galore! Accessed: 2025-09-08.

Langmead, B. and Salzberg, S. L. (2012). Fast gapped-read alignment with bowtie 2. Nat. Methods, 9(4):357–359.

Lasko, L. M., Jakob, C. G., Edalji, R. P., Qiu, W., Montgomery, D., Digiammarino, E. L., Hansen, T. M., Risi, R. M., Frey, R., Manaves, V., Shaw, B., Algire, M., Hessler, P., Lam, T., Uziel, T., Faivre, E., Ferguson, D., Buchanan, F. G., Martin, R. L., Torrent, M., Chiang, G. G., Karukurichi, K., Langston, J. W., Weinert, B. T., Choudhary, C., de Vries, P., Van Drie, J. H., McElligott, D., Kesicki, E., Marmorstein, R., Sun, C., Cole, P. A., Rosenberg, S. H., Michaelides, R., Lai, A., and Bromberg, K. D. (2017). Discovery of a selective catalytic p300/CBP inhibitor that targets lineage-specific tumours. Nature, 550(7674):128–132.

Lawrence, M., Huber, W., Pagès, H., Aboyoun, P., Carlson, M., Gentleman, R., Morgan, M. T., and Carey, V. J. (2013). Software for computing and annotating genomic ranges. PLoS Comput. Biol., 9(8):e1003118.

Li, H., Handsaker, B., Wysoker, A., Fennell, T., Ruan, J., Homer, N., Marth, G., Abecasis, G., Durbin, R., and 1000 Genome Project Data Processing Subgroup (2009). The sequence alignment/map format and SAMtools. Bioinformatics, 25(16):2078–2079.

Li, W., Notani, D., Ma, Q., Tanasa, B., Nunez, E., Chen, A. Y., Merkurjev, D., Zhang, J., Ohgi, K., Song, X., Oh, S., Kim, H.-S., Glass, C. K., and Rosenfeld, M. G. (2013). Functional roles of enhancer RNAs for oestrogen-dependent transcriptional activation. Nature, 498(7455):516–520.

Liu, Y., Shang, G., Zhang, X., Liu, F., Zhang, C., Li, Z., Jia, J., Xu, Y., Zhang, Z., Yang, S., Zhou, B., Luan, Y., Huang, Y., Peng, Y., Han, T., He, Y., and Zheng, H. (2022). CAMTA1 gene affects the ischemia-reperfusion injury by regulating CCND1. Front. Cell. Neurosci., 16:868291.

Lopez-Delisle, L., Rabbani, L., Wolff, J., Bhardwaj, V., Backofen, R., Grüning, B., Ramírez, F., and Manke, T. (2021). pyGenomeTracks: reproducible plots for multivariate genomic datasets. Bioinformatics, 37(3):422–423.

Lorenz, R., Bernhart, S. H., Höner Zu Siederdissen, C., Tafer, H., Flamm, C., Stadler, P. F., and Hofacker, I. L. (2011). ViennaRNA package 2.0. Algorithms Mol. Biol., 6:26.

Love, M. I., Huber, W., and Anders, S. (2014). Moderated estimation of fold change and dispersion for RNA-seq data with DESeq2. Genome Biol., 15(12):550.

Luo, Y., Hitz, B. C., Gabdank, I., Hilton, J. A., Kagda, M. S., Lam, B., Myers, Z., Sud, P., Jou, J., Lin, K., Baymuradov, U. K., Graham, K., Litton, C., Miyasato, S. R., Strattan, J. S., Jolanki, O., Lee, J.-W., Tanaka, F. Y., Adenekan, P., O’Neill, E., and Cherry, J. M. (2020). New developments on the encyclopedia of DNA elements (ENCODE) data portal. Nucleic Acids Res., 48(D1):D882–D889.

Manolio, T. A., Collins, F. S., Cox, N. J., Goldstein, D. B., Hindorff, L. A., Hunter, D. J., McCarthy, M. I., Ramos, E. M., Cardon, L. R., Chakravarti, A., Cho, J. H., Guttmacher, A. E., Kong, A., Kruglyak, L., Mardis, E., Rotimi, C. N., Slatkin, M., Valle, D., Whittemore, A. S., Boehnke, M., Clark, A. G., Eichler, E. E., Gibson, G., Haines, J. L., Mackay, T. F. C., McCarroll, S. A., and Visscher, P. M. (2009). Finding the missing heritability of complex diseases. Nature, 461(7265):747–753.

Mao, Z., Bonni, A., Xia, F., Nadal-Vicens, M., and Greenberg, M. E. (1999). Neuronal activity-dependent cell survival mediated by transcription factor MEF2. Science, 286(5440):785–790.

McCaskill, J. S. (1990). The equilibrium partition function and base pair binding probabilities for RNA secondary structure. Biopolymers, 29(6-7):1105–1119.

Miao, L., Tang, Y., Bonneau, A. R., Chan, S. H., Kojima, M. L., Pownall, M. E., Vejnar, C. E., Gao, F., Krishnaswamy, S., Hendry, C. E., and Giraldez, A. J. (2022). The landscape of pioneer factor activity reveals the mechanisms of chromatin reprogramming and genome activation. Mol. Cell, 82(5):986– 1002.e9.

Narita, T., Ito, S., Higashijima, Y., Chu, W. K., Neumann, K., Walter, J., Satpathy, S., Liebner, T., Hamilton, W. B., Maskey, E., Prus, G., Shibata, M., Iesmantavicius, V., Brickman, J. M., Anastassiadis, K., Koseki, H., and Choudhary, C. (2021). Enhancers are activated by p300/CBP activity-dependent PIC assembly, RNAPII recruitment, and pause release. Mol. Cell, 81(10):2166–2182.e6.

Ni, W.-J., Xie, F., and Leng, X.-M. (2020). Terminus-associated non-coding RNAs: Trash or treasure? Front. Genet., 11:552444.

Nora, E. P., Lajoie, B. R., Schulz, E. G., Giorgetti, L., Okamoto, I., Servant, N., Piolot, T., van Berkum, N. L., Meisig, J., Sedat, J., Gribnau, J., Barillot, E., Blüthgen, N., Dekker, J., and Heard, E. (2012). Spatial partitioning of the regulatory landscape of the X-inactivation centre. Nature, 485(7398):381– 385.

Parton, R. G. and del Pozo, M. A. (2013). Caveolae as plasma membrane sensors, protectors and organizers. Nat. Rev. Mol. Cell Biol., 14(2):98–112.

Pertea, G. and Pertea, M. (2020). GFF utilities: GffRead and GffCompare. F1000Research, 9(304):304.

Pertea, M., Pertea, G. M., Antonescu, C. M., Chang, T.-C., Mendell, J. T., and Salzberg, S. L. (2015). StringTie enables improved reconstruction of a transcriptome from RNA-seq reads. Nat. Biotechnol., 33(3):290–295.

Phillips-Cremins, J. E., Sauria, M. E. G., Sanyal, A., Gerasimova, T. I., Lajoie, B. R., Bell, J. S. K., Ong, C.-T., Hookway, T. A., Guo, C., Sun, Y., Bland, M. J., Wagstaff, W., Dalton, S., McDevitt, T. C., Sen, R., Dekker, J., Taylor, J., and Corces, V. G. (2013). Architectural protein subclasses shape 3D organization of genomes during lineage commitment. Cell, 153(6):1281–1295.

Pickrell, J. K. (2014). Joint analysis of functional genomic data and genome-wide association studies of 18 human traits. Am. J. Hum. Genet., 94(4):559–573.

Rada-Iglesias, A., Bajpai, R., Swigut, T., Brugmann, S. A., Flynn, R. A., and Wysocka, J. (2011). A unique chromatin signature uncovers early developmental enhancers in humans. Nature, 470(7333):279–283.

Ramírez, F., Bhardwaj, V., Arrigoni, L., Lam, K. C., Grüning, B. A., Villaveces, J., Habermann, B., Akhtar, A., and Manke, T. (2018). High-resolution TADs reveal DNA sequences underlying genome organization in flies. Nat. Commun., 9(1):189.

Ramírez, F., Ryan, D. P., Grüning, B., Bhardwaj, V., Kilpert, F., Richter, A. S., Heyne, S., Dündar, F., and Manke, T. (2016). deepTools2: a next generation web server for deep-sequencing data analysis. Nucleic Acids Res., 44(W1):W160–5.

Raney, B. J., Barber, G. P., Benet-Pagès, A., Casper, J., Clawson, H., Cline, M. S., Diekhans, M., Fischer, C., Navarro Gonzalez, J., Hickey, G., Hinrichs, A. S., Kuhn, R. M., Lee, B. T., Lee, C. M., Le Mercier, P., Miga, K. H., Nassar, L. R., Nejad, P., Paten, B., Perez, G., Schmelter, D., Speir, M. L., Wick, B. D., Zweig, A. S., Haussler, D., Kent, W. J., and Haeussler, M. (2024). The UCSC genome browser database: 2024 update. Nucleic Acids Res., 52(D1):D1082–D1088.

Rao, S. S. P., Huntley, M. H., Durand, N. C., Stamenova, E. K., Bochkov, I. D., Robinson, J. T., Sanborn, A. L., Machol, I., Omer, A. D., Lander, E. S., and Aiden, E. L. (2014). A 3D map of the human genome at kilobase resolution reveals principles of chromatin looping. Cell, 159(7):1665–1680.

Rauluseviciute, I., Riudavets-Puig, R., Blanc-Mathieu, R., Castro-Mondragon, J. A., Ferenc, K., Kumar, V., Lemma, R. B., Lucas, J., Chèneby, J., Baranasic, D., Khan, A., Fornes, O., Gundersen, S., Johansen, M., Hovig, E., Lenhard, B., Sandelin, A., Wasserman, W. W., Parcy, F., and Mathelier, A. (2024). JASPAR 2024: 20th anniversary of the open-access database of transcription factor binding profiles. Nucleic Acids Res., 52(D1):D174–D182.

Replogle, J. M., Bonnar, J. L., Pogson, A. N., Liem, C. R., Maier, K., Ding, Y., Russell, B. J., Wang, X., Leng, K., Guna, A., Norman, T. M., Pak, R. A., Ramos, D. M., Ward, M. E., Gilbert, L. A., Kampmann, M., Weissman, J. S., and Jost, M. (2022). Maximizing CRISPRi efficacy and accessibility with dual-sgRNA libraries and optimal effectors. Elife, 11.

Reuter, J. S. and Mathews, D. H. (2010). RNAstructure: software for RNA secondary structure prediction and analysis. BMC Bioinformatics, 11:129.

Reva, B., Antipin, Y., and Sander, C. (2011). Predicting the functional impact of protein mutations: application to cancer genomics. Nucleic Acids Res., 39(17):e118.

Rissone, P., Bizarro, C. V., and Ritort, F. (2022). Stem-loop formation drives RNA folding in mechanical unzipping experiments. Proc. Natl. Acad. Sci. U. S. A., 119(3):e2025575119.

Ritchie, M. E., Phipson, B., Wu, D., Hu, Y., Law, C. W., Shi, W., and Smyth, G. K. (2015). limma powers differential expression analyses for RNA-sequencing and microarray studies. Nucleic Acids Res., 43(7):e47.

Roadmap Epigenomics Consortium, Kundaje, A., Meuleman, W., Ernst, J., Bilenky, M., Yen, A., Heravi-Moussavi, A., Kheradpour, P., Zhang, Z., Wang, J., Ziller, M. J., Amin, V., Whitaker, J. W., Schultz, M. D., Ward, L. D., Sarkar, A., Quon, G., Sandstrom, R. S., Eaton, M. L., Wu, Y.-C., Pfenning, A. R., Wang, X., Claussnitzer, M., Liu, Y., Coarfa, C., Harris, R. A., Shoresh, N., Epstein, C. B., Gjoneska, E., Leung, D., Xie, W., Hawkins, R. D., Lister, R., Hong, C., Gascard, P., Mungall, A. J., Moore, R., Chuah, E., Tam, A., Canfield, T. K., Hansen, R. S., Kaul, R., Sabo, P. J., Bansal, M. S., Carles, A., Dixon, J. R., Farh, K.-H., Feizi, S., Karlic, R., Kim, A.-R., Kulkarni, A., Li, D., Lowdon, R., Elliott, G., Mercer, T. R., Neph, S. J., Onuchic, V., Polak, P., Rajagopal, N., Ray, P., Sallari, R. C., Siebenthall, K. T., Sinnott-Armstrong, N. A., Stevens, M., Thurman, R. E., Wu, J., Zhang, B., Zhou, X., Beaudet, A. E., Boyer, L. A., De Jager, P. L., Farnham, P. J., Fisher, S. J., Haussler, D., Jones, S. J. M., Li, W., Marra, M. A., McManus, M. T., Sunyaev, S., Thomson, J. A., Tlsty, T. D., Tsai, L.-H., Wang, W., Waterland, R. A., Zhang, M. Q., Chadwick, L. H., Bernstein, B. E., Costello, J. F., Ecker, J. R., Hirst, M., Meissner, A., Milosavljevic, A., Ren, B., Stamatoyannopoulos, J. A., Wang, T., and Kellis, M. (2015). Integrative analysis of 111 reference human epigenomes. Nature, 518(7539):317–330.

Ryan, M., Heverin, M., McLaughlin, R. L., and Hardiman, O. (2019). Lifetime risk and heritability of amyotrophic lateral sclerosis. JAMA Neurol., 76(11):1367–1374.

Sanjana, N. E., Shalem, O., and Zhang, F. (2014). Improved vectors and genome-wide libraries for CRISPR screening. Nat. Methods, 11(8):783–784.

Sawada, A., Wang, S., Jian, M., Leem, J., Wackerbarth, J., Egawa, J., Schilling, J. M., Platoshyn, O., Zemljic-Harpf, A., Roth, D. M., Patel, H. H., Patel, P. M., Marsala, M., and Head, B. P. (2019). Neuron-targeted caveolin-1 improves neuromuscular function and extends survival in SOD1G93A mice. FASEB J., 33(6):7545–7554.

Scherer, P. E., Lewis, R. Y., Volonte, D., Engelman, J. A., Galbiati, F., Couet, J., Kohtz, D. S., van Donselaar, E., Peters, P., and Lisanti, M. P. (1997). Cell-type and tissue-specific expression of caveolin-2. caveolins 1 and 2 co-localize and form a stable hetero-oligomeric complex in vivo. J. Biol. Chem., 272(46):29337–29346.

Schwalb, B., Michel, M., Zacher, B., Frühauf, K., Demel, C., Tresch, A., Gagneur, J., and Cramer, P. (2016). TT-seq maps the human transient transcriptome. Science, 352(6290):1225– 1228.

Seffens, W. and Digby, D. (1999). mRNAs have greater negative folding free energies than shuffled or codon choice randomized sequences. Nucleic Acids Res., 27(7):1578–1584.

Sexton, T., Yaffe, E., Kenigsberg, E., Bantignies, F., Leblanc, B., Hoichman, M., Parrinello, H., Tanay, A., and Cavalli, G. (2012). Three-dimensional folding and functional organization principles of the drosophila genome. Cell, 148(3):458–472.

Shalem, O., Sanjana, N. E., Hartenian, E., Shi, X., Scott, D. A., Mikkelson, T., Heckl, D., Ebert, B. L., Root, D. E., Doench, J. G., and Zhang, F. (2014). Genome-scale CRISPR-Cas9 knockout screening in human cells. Science, 343(6166):84–87.

Shalizi, A., Gaudillière, B., Yuan, Z., Stegmüller, J., Shirogane, T., Ge, Q., Tan, Y., Schulman, B., Harper, J. W., and Bonni, A. (2006). A calcium-regulated MEF2 sumoylation switch controls postsynaptic differentiation. Science, 311(5763):1012–1017.

Shen, Y., Yue, F., McCleary, D. F., Ye, Z., Edsall, L., Kuan, S., Wagner, U., Dixon, J., Lee, L., Lobanenkov, V. V., and Ren, B. (2012). A map of the cis-regulatory sequences in the mouse genome. Nature, 488(7409):116–120.

Shiraki, T., Kondo, S., Katayama, S., Waki, K., Kasukawa, T., Kawaji, H., Kodzius, R., Watahiki, A., Nakamura, M., Arakawa, T., Fukuda, S., Sasaki, D., Podhajska, A., Harbers, M., Kawai, J., Carninci, P., and Hayashizaki, Y. (2003). Cap analysis gene expression for high-throughput analysis of transcriptional starting point and identification of promoter usage. Proc. Natl. Acad. Sci. U. S. A., 100(26):15776–15781.

Siepel, A., Bejerano, G., Pedersen, J. S., Hinrichs, A. S., Hou, M., Rosenbloom, K., Clawson, H., Spieth, J., Hillier, L. W., Richards, S., Weinstock, G. M., Wilson, R. K., Gibbs, R. A., Kent, W. J., Miller, W., and Haussler, D. (2005). Evolutionarily conserved elements in vertebrate, insect, worm, and yeast genomes. Genome Res., 15(8):1034–1050.

Smith, T., Heger, A., and Sudbery, I. (2017). UMI-tools: modeling sequencing errors in unique molecular identifiers to improve quantification accuracy. Genome Res., 27(3):491–499.

Speed, D., Hemani, G., Johnson, M. R., and Balding, D. J. (2012). Improved heritability estimation from genome-wide SNPs. Am. J. Hum. Genet., 91(6):1011–1021.

Spitz, F. and Furlong, E. E. M. (2012). Transcription factors: from enhancer binding to developmental control. Nat. Rev. Genet., 13(9):613–626.

Svoboda, P. and Di Cara, A. (2006). Hairpin RNA: a secondary structure of primary importance. Cell. Mol. Life Sci., 63(7-8):901–908.

Takayasu, Y., Takeuchi, K., Kumari, R., Bennett, M. V. L., Zukin, R. S., and Francesconi, A. (2010). Caveolin-1 knockout mice exhibit impaired induction of mGluR-dependent long-term depression at CA3-CA1 synapses. Proc. Natl. Acad. Sci. U. S. A., 107(50):21778–21783.

Trotta, E. (2014). On the normalization of the minimum free energy of RNAs by sequence length. PLoS One, 9(11):e113380.

Trynka, G., Sandor, C., Han, B., Xu, H., Stranger, B. E., Liu, X. S., and Raychaudhuri, S. (2013). Chromatin marks identify critical cell types for fine mapping complex trait variants. Nat. Genet., 45(2):124–130.

van Arensbergen, J., Pagie, L., FitzPatrick, V. D., de Haas, M., Baltissen, M. P., Comoglio, F., van der Weide, R. H., Teunissen, H., Võsa, U., Franke, L., de Wit, E., Vermeulen, M., Bussemaker, H. J., and van Steensel, B. (2019). High-throughput identification of human SNPs affecting regulatory element activity. Nat. Genet., 51(7):1160–1169.

van Rheenen, W., Shatunov, A., Dekker, A. M., McLaughlin, R. L., Diekstra, F. P., Pulit, S. L., van der Spek, R. A. A., Võsa, U., de Jong, S., Robinson, M. R., Yang, J., Fogh, I., van Doormaal, P. T., Tazelaar, G. H. P., Koppers, M., Blokhuis, A. M., Sproviero, W., Jones, A. R., Kenna, K. P., van Eijk, K. R., Harschnitz, O., Schellevis, R. D., Brands, W. J., Medic, J., Menelaou, A., Vajda, A., Ticozzi, N., Lin, K., Rogelj, B., Vrabec, K., Ravnik-Glavač, M., Koritnik, B., Zidar, J., Leonardis, L., Grošelj, L. D., Millecamps, S., Salachas, F., Meininger, V., de Carvalho, M., Pinto, S., Mora, J. S., Rojas-García, R., Polak, M., Chandran, S., Colville, S., Swingler, R., Morrison, K. E., Shaw, P. J., Hardy, J., Orrell, R. W., Pittman, A., Sidle, K., Fratta, P., Malaspina, A., Topp, S., Petri, S., Abdulla, S., Drepper, C., Sendtner, M., Meyer, T., Ophoff, R. A., Staats, K. A., Wiedau-Pazos, M., Lomen-Hoerth, C., Van Deerlin, V. M., Trojanowski, J. Q., Elman, L., McCluskey, L., Basak, A. N., Tunca, C., Hamzeiy, H., Parman, Y., Meitinger, T., Lichtner, P., Radivojkov-Blagojevic, M., Andres, C. R., Maurel, C., Bensimon, G., Landwehrmeyer, B., Brice, A., Payan, C. A. M., Saker-Delye, S., Dürr, A., Wood, N. W., Tittmann, L., Lieb, W., Franke, A., Rietschel, M., Cichon, S., Nöthen, M. M., Amouyel, P., Tzourio, C., Dartigues, J.-F., Uitterlinden, A. G., Rivadeneira, F., Estrada, K., Hofman, A., Curtis, C., Blauw, H. M., van der Kooi, A. J., de Visser, M., Goris, A., Weber, M., Shaw, C. E., Smith, B. N., Pansarasa, O., Cereda, C., Del Bo, R., Comi, G. P., D’Alfonso, S., Bertolin, C., Sorarù, G., Mazzini, L., Pensato, V., Gellera, C., Tiloca, C., Ratti, A., Calvo, A., Moglia, C., Brunetti, M., Arcuti, S., Capozzo, R., Zecca, C., Lunetta, C., Penco, S., Riva, N., Padovani, A., Filosto, M., Muller, B., Stuit, R. J., PARALS Registry, SLALOM Group, SLAP Registry, FALS Sequencing Consortium, SLAGEN Consortium, NNIPPS Study Group, Blair, I., Zhang, K., McCann, E. P., Fifita, J. A., Nicholson, G. A., Rowe, D. B., Pamphlett, R., Kiernan, M. C., Grosskreutz, J., Witte, O. W., Ringer, T., Prell, T., Stubendorff, B., Kurth, I., Hübner, C. A., Leigh, P. N., Casale, F., Chio, A., Beghi, E., Pupillo, E., Tortelli, R., Logroscino, G., Powell, J., Ludolph, A. C., Weishaupt, J. H., Robberecht, W., Van Damme, P., Franke, L., Pers, T. H., Brown, R. H., Glass, J. D., Landers, J. E., Hardiman, O., Andersen, P. M., Corcia, P., Vourc’h, P., Silani, V., Wray, N. R., Visscher, P. M., de Bakker, P. I. W., van Es, M. A., Pasterkamp, R. J., Lewis, C. M., Breen, G., Al-Chalabi, A., van den Berg, L. H., and Veldink, J. H. (2016). Genome-wide association analyses identify new risk variants and the genetic architecture of amyotrophic lateral sclerosis. Nat. Genet., 48(9):1043–1048.

Watson, M., Park, Y., and Thoreen, C. (2020). Roadblock-qPCR: A simple and inexpensive strategy for targeted measurements of mRNA stability. RNA.

Welter, D., MacArthur, J., Morales, J., Burdett, T., Hall, P., Junkins, H., Klemm, A., Flicek, P., Manolio, T., Hindorff, L., and Parkinson, H. (2014). The NHGRI GWAS catalog, a curated resource of SNP-trait associations. Nucleic Acids Res., 42(Database issue):D1001–6.

Wickham, H. (2016). ggplot2: Elegant graphics for data analysis.

Xie, X., Lu, J., Kulbokas, E. J., Golub, T. R., Mootha, V., Lindblad-Toh, K., Lander, E. S., and Kellis, M. (2005). Systematic discovery of regulatory motifs in human promoters and 3’ UTRs by comparison of several mammals. Nature, 434(7031):338–345.

Yang, Q., She, H., Gearing, M., Colla, E., Lee, M., Shacka, J. J., and Mao, Z. (2009). Regulation of neuronal survival factor MEF2D by chaperone-mediated autophagy. Science, 323(5910):124–127.

Youn, H. D., Grozinger, C. M., and Liu, J. O. (2000). Calcium regulates transcriptional repression of myocyte enhancer factor 2 by histone deacetylase 4. J. Biol. Chem., 275(29):22563–22567.

Yousefian-Jazi, A., Sung, M. K., Lee, T., Hong, Y.-H., Choi, J. K., and Choi, J. (2020). Functional fine-mapping of noncoding risk variants in amyotrophic lateral sclerosis utilizing convolutional neural network. Sci. Rep., 10(1):12872.

Zhang, S., Cooper-Knock, J., Weimer, A. K., Shi, M., Moll, T., Marshall, J. N. G., Harvey, C., Nezhad, H. G., Franklin, J., Souza, C. D. S., Ning, K., Wang, C., Li, J., Dilliott, A. A., Farhan, S., Elhaik, E., Pasniceanu, I., Livesey, M. R., Eitan, C., Hornstein, E., Kenna, K. P., Project MinE ALS Sequencing Consortium, Veldink, J. H., Ferraiuolo, L., Shaw, P. J., and Snyder, M. P. (2022). Genome-wide identification of the genetic basis of amyotrophic lateral sclerosis. Neuron, 110(6):992–1008.e11.

Zhang, Y., Liu, T., Meyer, C. A., Eeckhoute, J., Johnson, D. S., Bernstein, B. E., Nusbaum, C., Myers, R. M., Brown, M., Li, W., and Liu, X. S. (2008). Model-based analysis of ChIP-seq (MACS). Genome Biol., 9(9):R137.

Zuin, J., Roth, G., Zhan, Y., Cramard, J., Redolfi, J., Piskadlo, E., Mach, P., Kryzhanovska, M., Tihanyi, G., Kohler, H., Eder, M., Leemans, C., van Steensel, B., Meister, P., Smallwood, S., and Giorgetti, L. (2022). Nonlinear control of transcription through enhancer-promoter interactions. Nature, 604(7906):571–577.

Zuker, M. and Stiegler, P. (1981). Optimal computer folding of large RNA sequences using thermodynamics and auxiliary information. Nucleic Acids Res., 9(1):133–148.

