## Supplementary figures and images for "Amyotrophic Lateral Sclerosis-associated 3′ UTR enhancer embedded within *CAV1* risk gene"

### Fig. S1: CTCF binds at TAD boundaries. CTCF binding as determined by ChIP-seq signal in SK-N-SH cells across all TADs genome-wide.

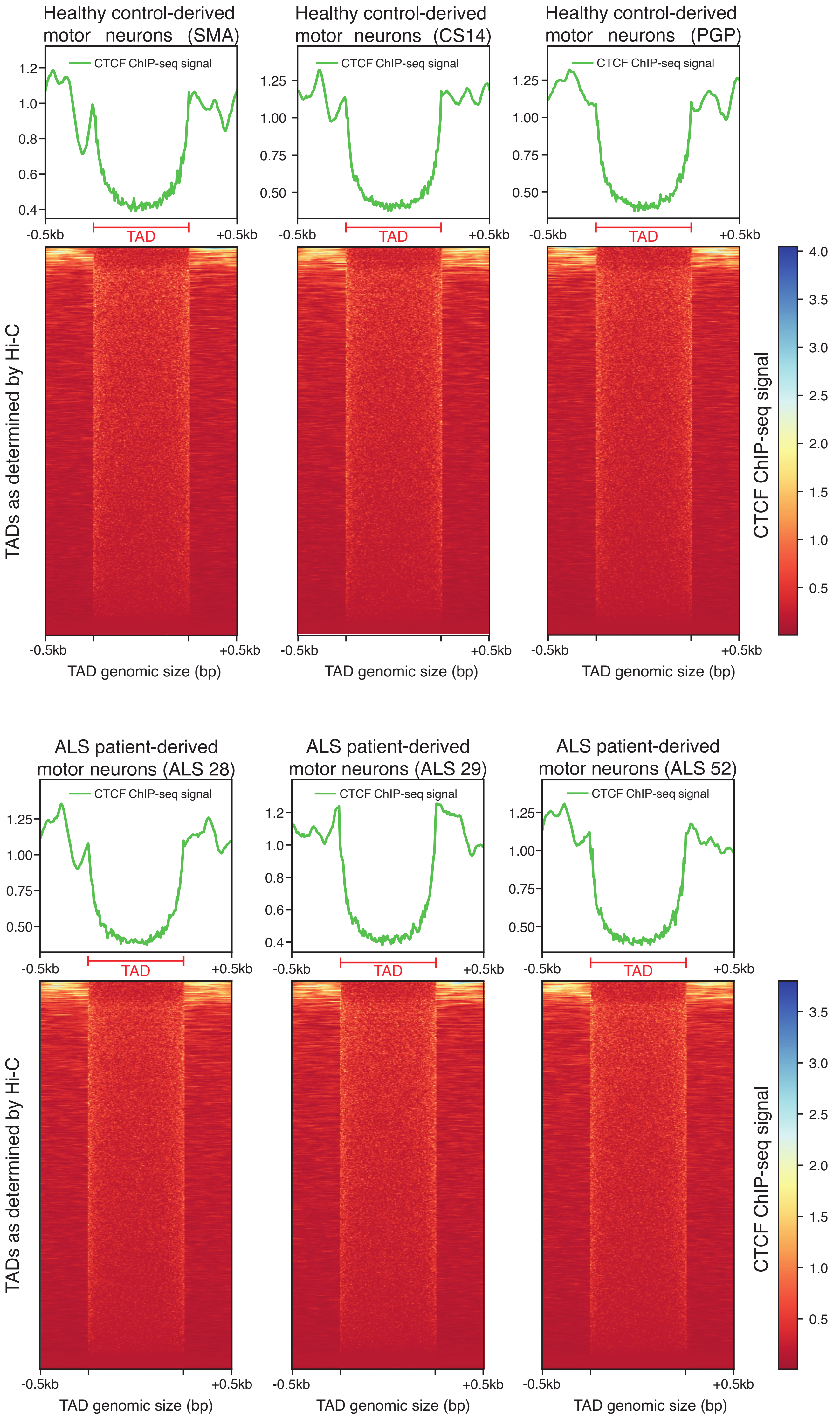

### Fig. S2: Residuals of the polynomial regression for MFEden versus GC-content.

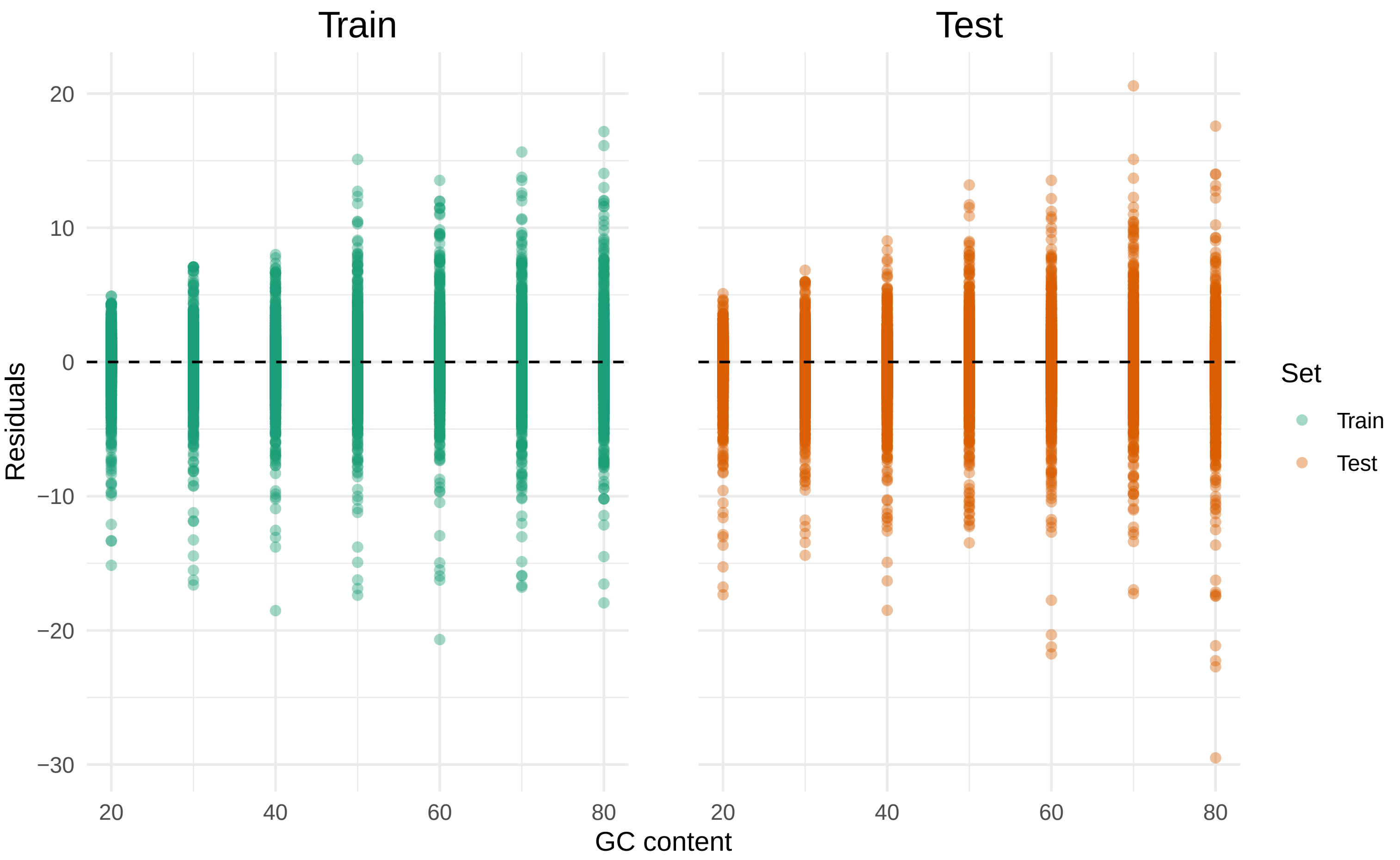
